# treeclimbR pinpoints the data-dependent resolution of hierarchical hypotheses

**DOI:** 10.1101/2020.06.08.140608

**Authors:** Ruizhu Huang, Charlotte Soneson, Pierre-Luc Germain, Thomas S.B. Schmidt, Christian Von Mering, Mark D. Robinson

## Abstract

The arrangement of hypotheses in a hierarchical structure (e.g., phylogenies, cell types) appears in many research fields and indicates different resolutions at which data can be interpreted. A common goal is to find a representative resolution that gives high sensitivity to identify relevant entities (e.g., microbial taxa or cell subpopulations) that are related to a phenotypic outcome (e.g. disease status) while controlling false detections, therefore providing a more compact view of detected entities and summarizing characteristics shared among them. Current methods, either performing hypothesis tests at an arbitrary resolution or testing hypotheses at all possible resolutions leading to nested results, are suboptimal. Moreover, they are not flexible enough to work in situations where each entity has multiple features to consider and different resolutions might be required for different features. For example, in single cell RNA-seq data, an increasing focus is to find differential state genes that change expression within a cell subpopulation in response to an external stimulus. Such differential expression might occur at different resolutions (e.g., all cells or a small set of cells) for different genes. Our new algorithm *treeclimbR* is designed to fill this gap by exploiting a hierarchical tree of entities, proposing multiple candidates that capture the latent signal and pinpointing branches or leaves that contain features of interest, in a data-driven way. It outperforms currently available methods on synthetic data, and we highlight the approach on various applications, including microbiome and microRNA surveys as well as single cell cytometry and RNA-seq datasets. With the emergence of various multi-resolution genomic datasets, *treeclimbR* provides a thorough inspection on entities across resolutions and gives additional flexibility to uncover biological associations.

## Introduction

In many fields, multiple hypotheses are simultaneously tested to investigate the association between a phenotypic outcome (e.g., disease status) and measured entities (e.g., microbial taxa). When a hierarchy of entities exists, hypotheses can be arranged in a tree structure that indicates different resolutions of interpretation. For example, in metagenomics, a tree constructed based on marker gene (or genomic) sequence provides taxonomic resolution to investigate associations between phenotype and taxa abundance. Associations tested only at a fine resolution (e.g., species-level on the taxonomic tree) might not have sufficient statistical power to detect taxa with small changes, which are of interest if they appear coherently. Given that closely related taxa often share similarity in response to environmental change [1], differential analysis performed on a broader resolution (e.g., phylum level) may improve detection by accumulating the small coherent changes. However, a broad resolution is not always desirable: using too low of a resolution cannot pinpoint specific taxa that exhibit an association. Thus, there is a need for methods that balance detection power and error control, while also giving flexibility to find the relevant resolution to interpret the data.

A similar challenge exists in the analysis of microRNA (miRNA) data. MiRNAs are small non-coding RNA molecules, and their dysregulation is associated with diseases including retinal disorder, cardiovascular disease, and cancer [2, 3]. The abundance of miRNAs could be affected by regulation occuring at multiple levels of their biogenesis [4]: miRNAs in the same transcript are generally co-transcribed but the individual miRNAs can be additionally regulated at the post transcriptional level, and variations in Dicer cleavage or RNA editing can lead to distinct RNA fragments. A tree, where each leaf represents a unique mature miRNA sequence, and internal nodes represent miRNA duplexes, primary transcript and clusters of miRNAs, could provide different biogenesis resolutions to interpret disease-associated miRNA dysregulation. A typical approach for microbial and miRNA surveys is so-called differential abundance analysis (DA) [5, 6], where the abundance of each entity measured is tested for association with a phenotype of interest. Regardless of the specific data, the focus becomes locating the right resolution (e.g., microbial taxa or miRNAs) that have phenotype-associated abundance changes. In such cases, the input data shares the same structure: abundance of entities collected across samples and a tree encoding the hierarchy of entities. A similar but more complicated case is so-called differential state analysis (DS) [7, 8], which arises in the analysis of single cell datasets and typically involves comparing measurements on a single entity (e.g., cell subpopulation) across multiple samples (e.g., changes in marker intensity amongst markers not used in subpopulation definition). In contrast to DA tests, DS can have considerable multiplicity, with 10s to 1000s of feature profiles (e.g., antibody intensity or gene expression) for each entity. Such scenarios are usually encountered in single cell RNA sequencing (scRNAseq) data and mass cytometry (CyTOF) data, where several reports have shown subpopulation-specific responses that occur in disease states or due to external stimuli [9, 10, 11, 12]. Notably, the classification of cell subpopulations often requires selecting a resolution of the data, and even when well-established markers exist, a cell subpopulation might still contain hidden diversity [13, 14]. It is also unclear whether detected state changes really occur at the subpopulation level or are driven by smaller subsets of cells. In the extreme case, if changes occur at a fine resolution and in offsetting directions, they might not even be detected when the whole subpopulation is considered. It is therefore desirable to have more flexibility in the analysis, where some changes of interest occur at very specific subpopulations, while others occur amongst broad cell subpopulations. To achieve this, the use of a tree to store cell subpopulations on different resolutions, and exploring on the tree to find a suitable resolution, will ideally lead to better understanding of cellular response. Briefly, in the DS test, data includes a tree encoding the hierarchy of entities (cell subpopulations), and observations of multiple features (genes or antibodies) on each entity across samples. Notably, the DA test is a special case of the DS test, where each entity has only one feature: relative abundance.

Currently, several methods are available, either general for multiplicity correction or specific for a certain type of data. Yekutieli [15] proposed the hierarchical false discovery rate controlling procedure (*HFDR*) for tree-structured hypotheses. It increases power by selectively focusing on branches that are more likely to contain alternative hypotheses. Instead of generating hierarchical hypotheses, an empirical Bayes approach, *StructFDR* [16], performs hypothesis tests only on the leaf level and improves the power by incorporating a correlation matrix converted from a tree (based on distances among leaves) as the prior correlation structure to share information among hypotheses. *MiLineage* [17] is developed for microbiome data and localizes the phenotype-associated lineages on the taxonomic tree by splitting a tree into multiple lineages, each of which includes a parent node (taxon) and its direct child nodes (taxa on a finer resolution). It then performs multivariate tests concerning multiple taxa in a lineage to test the association of lineage to a phenotypic outcome. *LEfSe* [18] mainly focuses on biomarker discovery of metagenomic data, by first identifying DA features using the Kruskal-Wallis sum-rank test (KW), and further selects features that have effect sizes above a specified threshold using linear discriminant analysis. *Citrus* [19] works on CyTOF data and applies a lasso-regularized regression model [20] to automatically select stratifying subpopulations and cell response features that are the best predictors of a phenotypic outcome. An alternative to *Citrus* [19], *diffcyt* [6], over-clusters cells into subpopulations and performs differential analysis at this higher resolution separately for each feature, without any attempt to summarize concordant signal on similar cell subpopulations.

Existing methods have limitations. *HFDR* [15] does not perform well for compositional data in the DA setting because it typically stops right on the root branch, where essentially sample-level sequencing depths are compared and thus it fails to move along branches to specific entities; furthermore, no specific consideration is given to the DS case where there are multiple hypotheses (multiple features) per node: the global FDR over all features cannot be controlled at a specific level if the procedure is performed separately on each feature, and decisions of rejecting a node to move toward its child nodes cannot be taken separately for different features if the procedure is performed simultaneously on all features. *StructFDR* [16], which transforms P-values into z-scores and performs z-score smoothing among leaves in close proximity (leveraging the tree structure), is powerful to identify clustered signals. However, when signals are scattered in the tree, their z-scores might be pulled down by their non-signal neighbors due to smoothing, which makes *StructFDR* less powerful than BH [21], as shown by Bichat *et al.* [22]; additionally, no consideration is made for the DS case where a leaf has multiple P-values. *LEfSe* [18] directly applies the KW test on each feature and thus does not take confounders into consideration, and might have much higher FDR than expected due to the lack of multiplicity correction. Lasso-regularized models [20] (e.g., *Citrus* [19]), which tend to pick one and ignore the rest among highly correlated predictors [23], can be potentially applied to pick a resolution of a relevant branch where nodes representing a cell subpopulation are nested and highly correlated. However, the automatic selection might also occur among highly correlated cell subpopulations from different branches, or features (e.g., genes) behaving similarly in the same cell subpopulations, which leads to loss of relevant information. *diffcyt* works well for the DA and DS case of CyToF data but at a fixed arbitrary resolution.

To overcome these limitations, we propose a new algorithm, *treeclimbR*, that uses the tree topology together with the molecular profiling data. We show gains in sensitivity to detect relevant entities when a tree has branches with coherent changes, and similar performance to BH [21] when the tree is uninformative. *treeclimbR* has several unique attributes: it explores the latent resolution of association by proposing multiple candidate resolutions and it selects the optimal candidate in a data-driven way; since each candidate resolution consists of nodes that do not have ancestor-descendant relationship, *treeclimbR* can identify branches of relevant entities to show characteristics shared among them while avoiding nested nodes that are difficult to interpret. Furthermore, in DS testing, the exploration of resolution is conducted separately for each feature, which allows different features to stop at different resolutions of the tree. This matches the reality that features (e.g., gene expression) might be regulated differently in different cell subpopulations and therefore allows a more flexible data analysis platform.

## Results

### Overview of treeclimbR

There are many examples in the biology of entities (such as miRNAs, microbial species or single cell populations), measured across samples from different conditions, where it is of interest to detect associations between (the presence of) entities and condition (e.g., disease status), and where information of the relationship *between* entities could be leveraged to simplify or bring insight into the interpretation. Our new algorithm, *treeclimbR*, combines a tree that encodes the hierarchical relationship between entities with these observations, and pinpoints a suitable resolution on the tree to interpret the association. In this manuscript, branches of relevant entities that have differential abundance or expression levels among groups are called signal branches. Notably, the approach is general, in that it can be applied to interpret arbitrary statistical models that are fit at the nodes of the tree. The five main steps are illustrated in Figure 1: (a) data aggregation, (b) differential analysis, (c) candidate proposal, (d) multiple testing correction, and (e) candidate evaluation.

**Figure 1:**
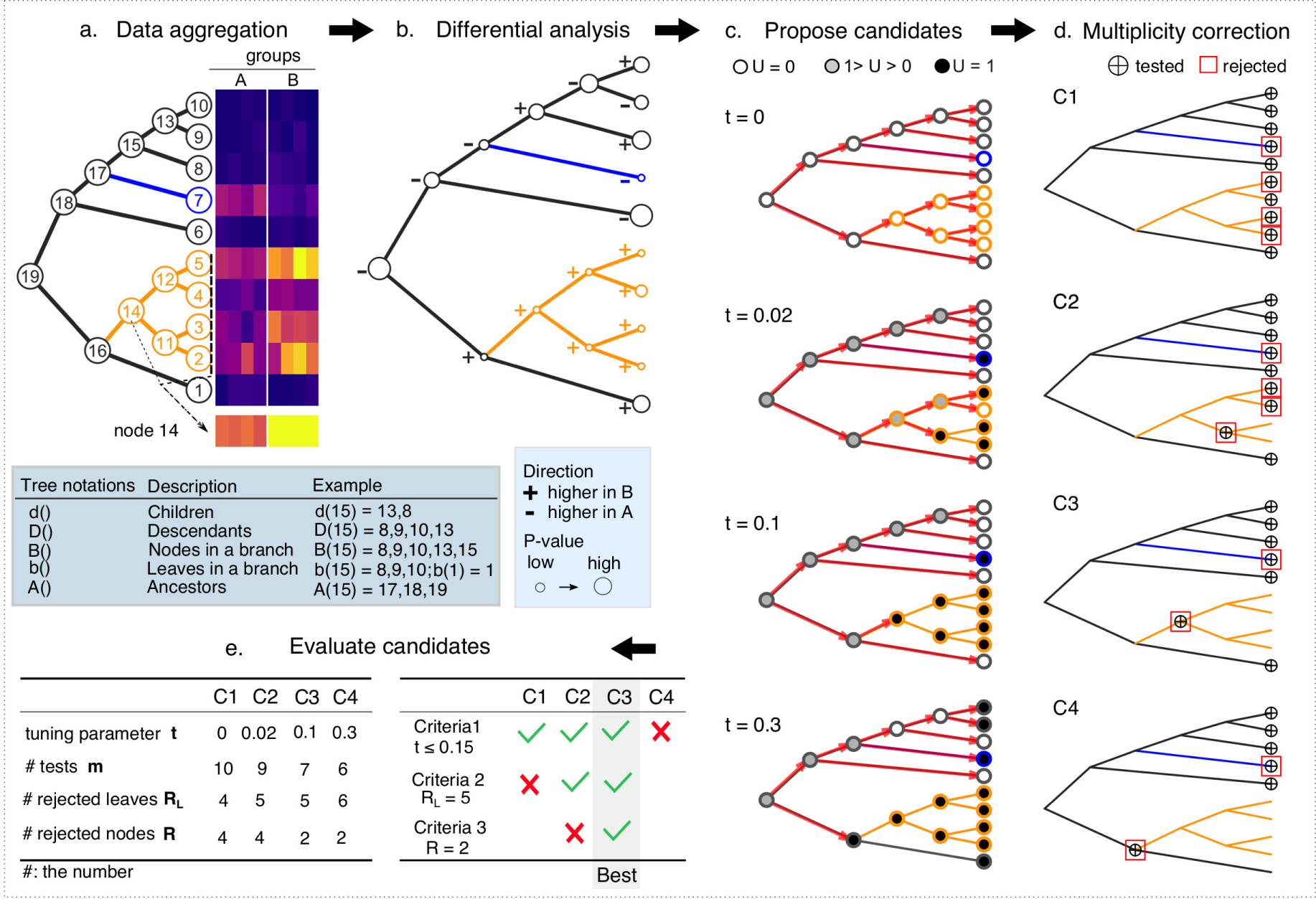
Schematic overview of the treeclimbR algorithm. **(a)** Data aggregation. An example tree of entities with 10 leaves (1 *−* 10) and 9 internal nodes (11 *−* 19). Measurements of entities in samples across multiple groups (e.g., A and B) are shown in the heatmap. For internal nodes, data is generated from their descendant leaves (e.g., node 14 from nodes 2, 3, 4, 5). Signal branches are colored in blue (higher in A) and orange (higher in B). **(b)** Differential analysis is run on each node to estimate its direction of change and obtain a P-value. **(c)** The generation of four example candidates, *C*1, *C*2, *C*3 and *C*4. The climbing starts from the root and stops when reaching leaves or nodes with *U* = 1, a score dependent on a tuning parameter *t*. **(d)** Multiple testing correction on each candidate. Nodes labelled by are ⊕ termini (i.e., candidates) obtained in **(c)**. *C*1 is the leaf level where differential analysis is performed when a tree is not available. The null hypothesis is tested on each node of a candidate, and multiplicity is corrected within a candidate. Rejected nodes are shown in red rectangles. **(e)** Results from **(c)** are summarized in the left table. *m, R* and *R*_*L*_ are the number of hypothesis tests (nodes with ⊕), the number of rejected nodes, and the number of rejected leaves (the descendant leaves of rejected nodes), respectively. The best candidate is selected based on three criteria. Candidates that fail in one criterion would not enter to the next. **Note:** Tree notations used in this article are listed below **(a)**.

The data aggregation and differential analysis generate data and statistics for internal nodes. Depending on the context, for each internal node, we either take the mean, median or sum of the data within its descendant leaves. On each node, we compare (aggregated) data across groups to get an estimated direction of change, and test a null hypothesis, *H*_0_ (e.g., that there is no difference), resulting in a P-value. As shown in Figure 1b, the hypotheses are in a hierarchical structure that might affect the control of false discovery rate (FDR) when using methods (e.g., the Benjaminin-Hochberg procedure [21]) to correct for multiplicity. To solve the hierarchical issue, an internal node is used to represent its descendant leaves that have coherent change. As the true signal is unknown, we explore the whole tree using a search procedure that starts from the root, and moves toward the leaves to capture the latent signal pattern at different resolutions, which we refer to as “candidates”.

Multiple candidates are proposed and a selection process is applied to select the optimal one. Figure 1c shows the generation of four example candidates (*C1, C2, C3, C4*) based on node-level *U* scores, which combine the direction and strength of the association and vary with a parameter *t* that has range [0, 1] (see Methods). The whole tree can be scanned and the search is stopped at different granularities (labelled as ‘tested’ in Figure 1d) to propose multiple candidates. If the null hypothesis on an internal node is rejected, all its descendant leaves are considered to have their null hypotheses rejected. Multiple hypothesis correction is performed separately on each candidate (see Figure 1e). The best candidate is selected by evaluating candidates according to three criteria: i) restricting the range of *t* (to control the FDR on the leaf level), which is determined by the average size of signal branches that could be detected, and is therefore data-dependent (see Methods); ii) selecting candidates with more rejected leaf nodes to increase the power to detect entities with signal; iii) selecting the signal branches with fewest internal nodes (e.g. C3 over C2 in Figure 1d), which makes the interpretation easier, and is desirable to find the right resolution.

Importantly, the procedure described in Figure 1 is for the DA test, where each entity has one feature (i.e., relative abundance across samples). A similar overall procedure is applied to the DS case, except that each entity has multiple features (e.g., multiple markers or genes) and the features are allowed to have associations at different resolutions; the candidate selection process is thus repeated for each feature separately.

### Performance assessment on synthetic datasets

We demonstrate the performance of *treeclimbR* against several competing methods, including *miLineage* [17], *StructFDR* [16], *HFDR* [15], *BH [21], minP* (see Supplementary Note 3), *LefSe* [18] and lasso-regularized logistic regression (*lasso*) [23] on synthetic microbial datasets (parametric and non-parametric), and two published semi-simulated single cell mass cytometry (CyTOF) datasets [6] (*AML-sim* and *BCR-XL-sim*).

#### Parametric synthetic microbial datasets

Operational taxonomic unit (OTU) counts are sampled from Dirichlet-multinomial distributions based on the real data in three scenarios adapted from Xiao *et al.* [16] *(see Methods): balanced signal (BS*), unbalanced signal (*US*), sporadic signal (*SS*) as shown schematically in Figure 2a. Instead of directly multiplying counts of selected OTUs in the treatment group by a fold change, we simulate differences by modifying parameters to introduce DA (see Methods). This ensures that relative abundance of non-DA OTUs remains fixed between groups in the compositional data, and better simulates differences of low-abundance OTUs that might otherwise have zero counts. Each scenario has two signal branches where OTUs have DA between the control and treatment group; OTUs in the same signal branch change in the same direction. In *BS*, the fold changes of OTUs within the signal branch are fixed, whereas the fold changes in the *US* case are (in the same direction but) different in magnitude. *SS* is similar to *BS*, except that only subsets of OTUs change (the rest remain unchanged). The library size is randomly drawn from library sizes of the 153 samples in the *throat_v35* data set (see Methods). For each scenario, we simulate different sample sizes: 10, 25 and 50 per group. In each combination of scenario and sample size, 5 repetitions are made.

**Figure 2:**
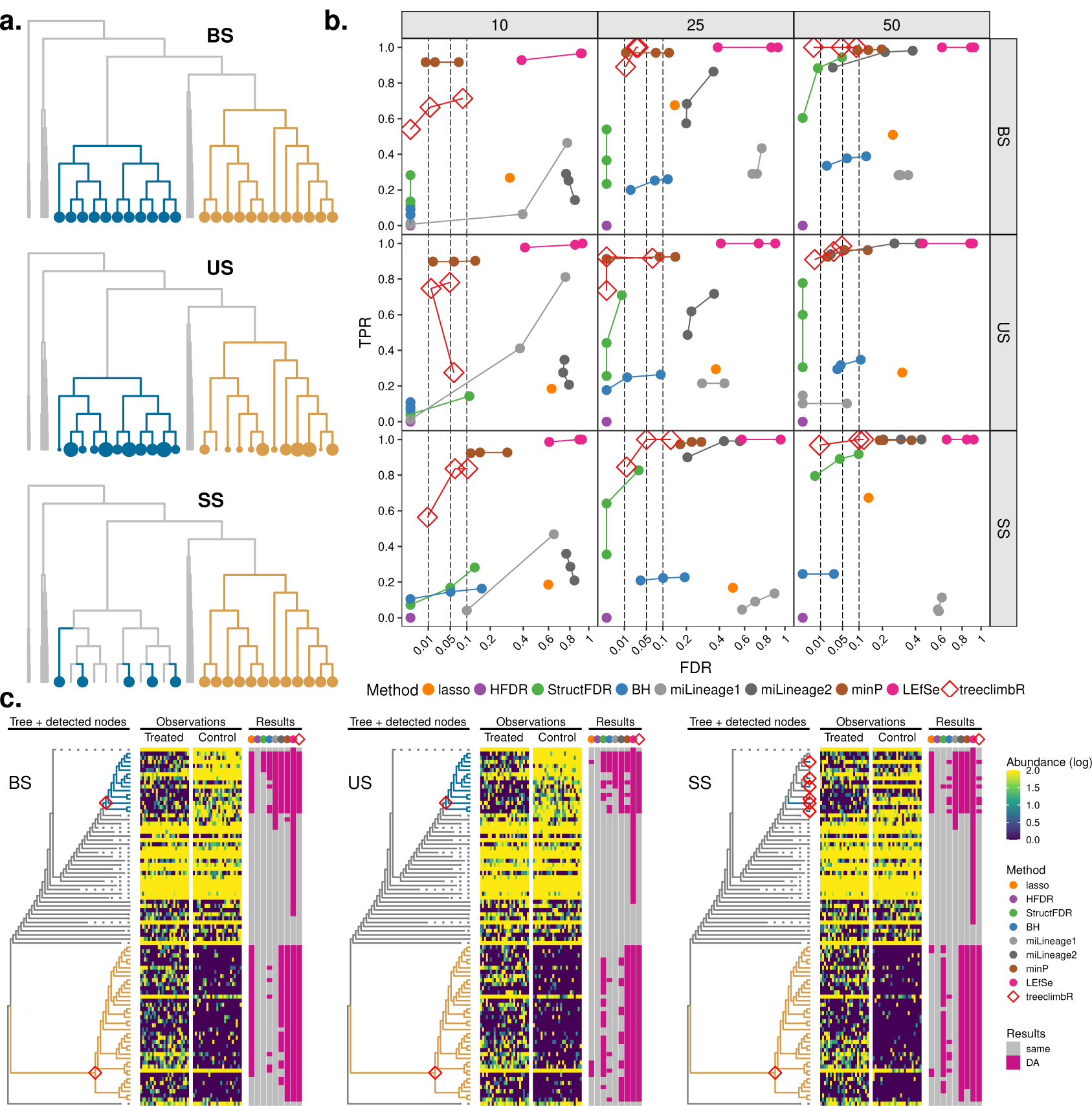
The performance of methods on parametric simulated microbial data. **(a)**. A schematic example of three simulated scenarios: BS, US and SS. Signal branches (i.e., with DA between groups) are in turquoise (decreased) and gold (increased), and larger points represent bigger change. **(b)**. Average TPR and FDR (over 5 repetitions) of methods on three scenarios under different sample sizes (10, 25, and 50 per group). Methods are in colors. Each method has three points that represent imposed FDR cutoffs at 0.01, 0.05 and 0.1. **(c)**. DA branches identified by methods in one of the 5 repetitions. The non-DA branches of the phylogenetic tree are represented with dashed lines to save space. Nodes identified by *treeclimbR* are labeled on the tree; and other methods are in Supplementary Figure 1. OTU counts are shown in the heatmap with samples split by groups. All OTUs (rows of heatmap) identified as DA by methods are summarized in the results panel.

The average performance of 5 repeated simulations is shown in Figure 2b. Both *lasso* and *miLineage* identify nested nodes and cannot pinpoint DA branches. If identified nodes that are closest to the root are used, OTUs reported by *miLineage* and *lasso* are mostly false positives (see Supplementary Figure 2). Here, to minimize the FDR of *lasso* and *miLineage*, we use their identified nodes that are closest to the leaf level. Generally, methods using a tree, such as *treeclimbR, StructFDR* [16], *and minP*, have higher power than *BH* [21]. *HFDR* [15] is unable to detect any changes between groups, because it starts the search from the root of the tree, which effectively represents the sequencing depth of samples, and typically stops right at the root where the null hypothesis cannot be rejected; thus, its TPR and FDR are equal to zero. In all scenarios, *treeclimbR* outperforms others with high TPR and well-controlled FDR. *minP* performs well with high TPR in all scenarios, but does not always control the FDR in the *SS* scenario where the signal does not occupy a full branch. In all three scenarios, *lasso* [23] *and miLineage* [17] have much higher FDR than expected. At a 5% FDR cutoff, OTUs identified by methods on three simulated scenarios with 25 samples per group are compared in Figure 2c. *BH* [21] fails to find some OTUs due to low abundance or low fold change. *treeclimbR* manages to aggregate concordant signal and to stop at the right level of the tree. The two-part analysis of *miLineage* (*miLineage2*) manages to detect some OTUs with sparse counts in the gold branch, while the one-part analysis (*miLineage1*) does not. *LEfSe* [18] identifies almost all DA branches in all simulations but with a lot of false discoveries, which may be due to the lack of multiplicity correction.

#### Non-parametric synthetic microbial datasets

Recently, Bichat *et al.* [22] have shown that currently available tree-based procedures, *StructFDR* [16] *or HFDR* [15] do not outperform the classical BH procedure when analyzing microbial data organized onto a taxonomic or phylogenetic tree. Even worse, they show that tree-based procedures might have a negative effect, giving either lower power or slightly higher power but poor FDR control compared to BH [21]. Their simulation is based on a real microbial dataset. Notably, they simulate differences by randomly selecting a set of OTUs from the most prevalent ones, and multiply counts in one of the experimental conditions by a fold change (e.g., 5). In other words, they simulate a tree that is uninformative, which provides us a negative control; we have reproduced their results and add the performance of *treeclimbR* to their non-parametric simulation (see Supplementary Figure 3). Tree-based methods offer no advantage when the tree is uninformative, whereas *treeclimbR* performs in this case on par with BH [21], in terms of both power and error control.

#### AML-sim

We next use a dataset that simulates the phenotype of minimal residual disease in acute myeloid leukemia (AML) patients, which is designed to evaluate the performance of DA methods after clustering CyTOF profiles according to a set of lineage markers. The *AML-sim* dataset provides simulations for two subtypes of AML (cytogenetically normal, CN; and core-binding factor translocation, CBF); only results on subtype CN are shown. The data consists of 5 healthy and 5 synthetic “diseased” samples that are generated by spiking in a small percentage of AML blast cells from CN samples into healthy samples [6]. AML blast cells are sufficiently distinct and can typically be clustered into a separate subpopulation. Depending on the proportion of spiked-in cells, the simulated scenarios are considered as strong (5%), medium (1%) and weak (0.1%).

We follow the concept of *diffcyt* [6] to group cells into a large number of clusters using the *FlowSOM* algorithm, and compare the abundance of clusters between the healthy and diseased groups for each cluster. Here, three different numbers of clusters have been tried: 400, 900 and 1600. A tree is built from the generated clusters based on the median expression of lineage markers (see Methods). TPR-FDR performances are shown in Figure 3a and a summary of each method’s detections in the context of related cells is shown in Figure 3b. *HFDR* is unable to detect the simulated signal, and has TPR and FDR both equal to 0 in all scenarios. Other methods perform well with high TPR and low FDR in the medium and the strong scenarios, and all methods fail to detect the weak signal. In the medium scenario, *diffcyt*’s TPR drops slightly when a large number of clusters (e.g., 1600) is used. For the *medium* scenario with 900 clusters, *treeclimbR, minP* and *diffcyt* detect the same branch that mainly includes the AML blast cells from CN samples: *treeclimbR* and *minP* reveal an internal node, and *diffcyt* highlights the three descendant leaves of the internal node. *StructFDR* misses one leaf that contains mostly AML cells. Compared to *treeclimbR, lasso* identifies an additional leaf that contains mostly non-AML cells.

**Figure 3:**
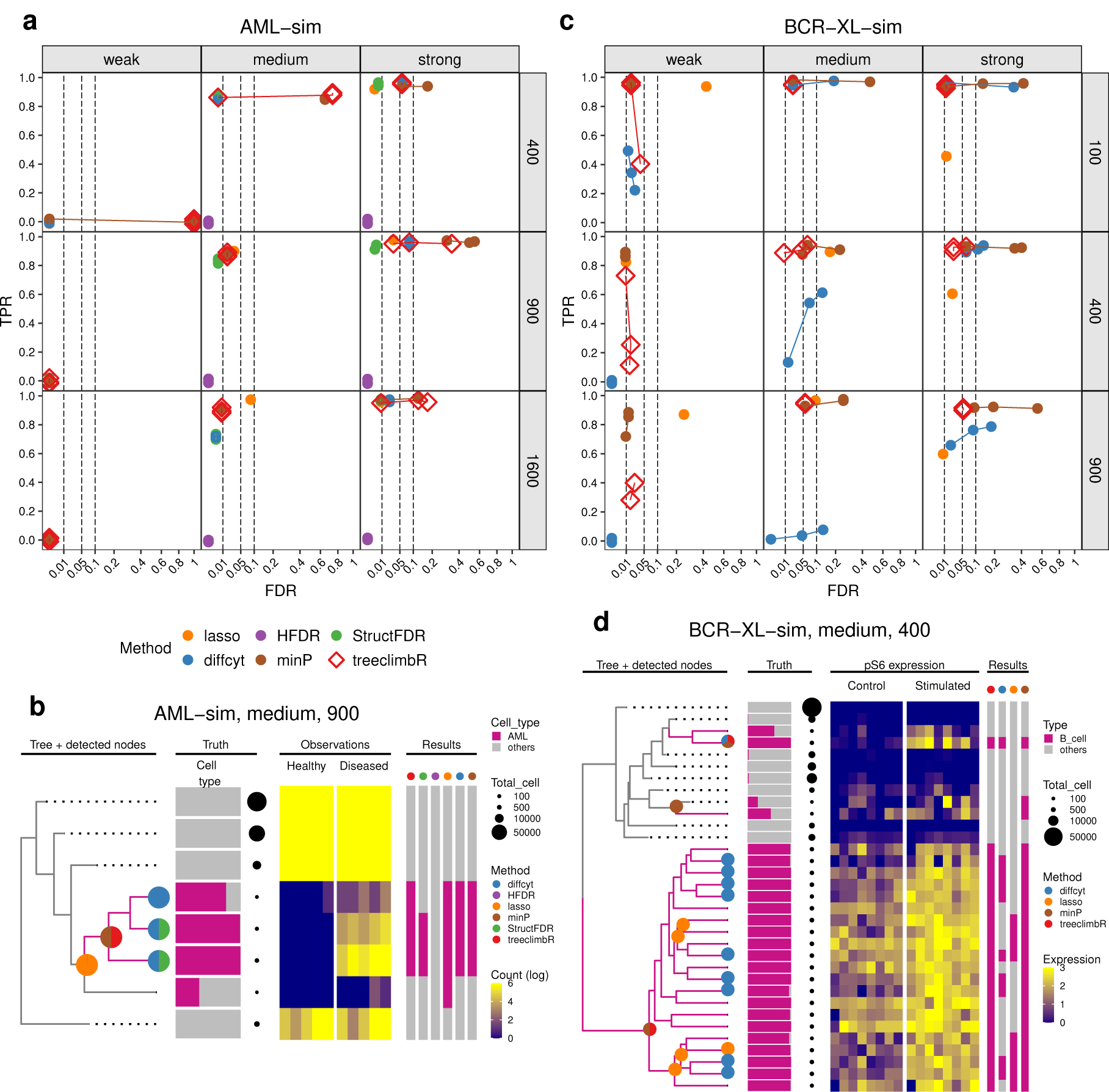
Results on two semi-simulated datasets (AML-sim and BCR-XL-sim). **(a)** TPR vs. FDR of methods on *AML-sim* under three scenarios (weak, medium, and strong) when trees have 400, 900 and 1600 leaves. Methods are in colors. FDR cutoffs are at 0.01, 0.05 and 0.1. **(b)** DA branches identified by methods in the medium scenario of *AML-sim* using a tree with 900 leaves. The four panels show the tree, truth, observations and results. The tree panel displays non-DA branches (without AML blast cells) in dashed lines to save space, colors branches with AML blast cells above 50% in purple. and labels nodes identified by methods; the truth panel shows cell type compositions and cell counts (point sizes) of leaves; the observation panel shows cell counts on leaves (rows) in samples (columns) split by groups; the result panel annotates leaves identified by methods. **(c)** same as **(a)**, except the *BCR-XL-sim* dataset is presented. **(d)** same as **(b)**, except the medium scenario of *BCR-XL-sim* using 400-leaf tree is presented. Here, only one feature (*pS6*) is shown.

#### BCR-XL-sim

We next test a dataset that consists of 8 paired samples of peripheral blood mononuclear cells (PBMCs) in two treatment groups: untreated and stimulated with B cell receptor*/*Fc receptor cross linker (BCR-XL); the goal is to detect DS within subpopulations. Samples in the control group have healthy PBMCs, and those in stimulated group are simulated from healthy PBMCs with spiked-in B cells from BCR-XL stimulated samples [6]. In other words, samples in the two groups are different in the expression of some protein markers, including pS6, pPlcg2, pErk, and pNFkB, in B cells. The difference in marker expression profiles between two groups is scaled to make groups distinct at three different levels: weak, medium and strong. Cells are again grouped into a large number of clusters using *FlowSOM*, and the expression of a protein marker on each cluster is compared between the control and the stimulated groups. Three numbers of clusters have been used: 100, 400 and 900. The tree is again built using the median expression of lineage markers in clusters.

TPR and FDR performance is calculated at the cell level, as shown in Figure 3b. A true positive is a (spiked-in) B cell found in a DS cluster that has at least one protein marker identified as differentially expressed between groups, and a false positive is a non-B-cell found in a cluster deemed DS. For the medium and strong scenarios, *treeclimbR* performs well with high TPR and controlled FDR; *minP* shows results similar to *treeclimbR* but with higher FDR; *diffcyt* works well with 100 clusters, but its TPR decreases as the number of clusters increases; *lasso* has slightly lower TPR than *treeclimbR* and *minP*. Signal branches identified in the medium scenario using 400 clusters are shown in Figure 3d for a single marker protein, *pS6*. Both *treeclimbR* and *minP* identify a large branch of B cells by picking its branch node, while *diffcyt* and *lasso* find only some of its leaves or sub-branches. In Figure 3b, *lasso* displays almost equal TPR as *treeclimbR* because most of those missing sub-branches are identified in other marker proteins (see Supplementary Figure 4). Because of the selection that *lasso* models apply, it might fail to identify some DS clusters for individual protein markers that are highly correlated with other strongly-associated markers. Additionally, the result of *lasso* includes nested nodes, which can be difficult to interpret.

### Tree-assisted DA and DS analyses

To highlight the diversity of applications where tree-assisted DA or DS detection arises, we applied *treeclimbR* to three datasets, including gut microbiota data, mouse miRNA data and mouse cortex scRNAseq data.

#### Differential abundance of microbes in infants born differently

We applied *treeclimbR* to a public metagenomic shotgun sequencing study on fecal samples [24], with the aim to investigate whether babies born vaginally or by C-section have different microbiome compositions (see Methods). The dataset includes 464 metaOTUs and a total of 381 samples collected across the two groups at different time points: 4 days (0M), 4 months (4M) and 12 months (12M), as shown in Figure 4. Nodes reported as DA by *treeclimbR* are according to a 5% FDR cutoff. In particular, at 0M, 8 branches and 7 leaves (in total, 188 metaOTUs) are detected to be DA between C-section and vaginal babies; the difference becomes less distinct as babies grow: 2 branches and 5 leaves (65 metaOTUs) and 8 leaves are detected at 4M and 12M, respectively. The main change in composition comes from the *Bacteroides* genus, which was previously shown to be less abundant in C-section babies [25]. Vaginal babies are enriched for species in genera (e.g., *Prevotella* and *Lactobacillus*) that resemble their mother’s vaginal microbiota, whereas C-section newborns tend to have higher abundance of species in genera (e.g., *Staphylococcus*) that are likely to be acquired from the hospital environment or from the mother’s skin [26].

**Figure 4:**
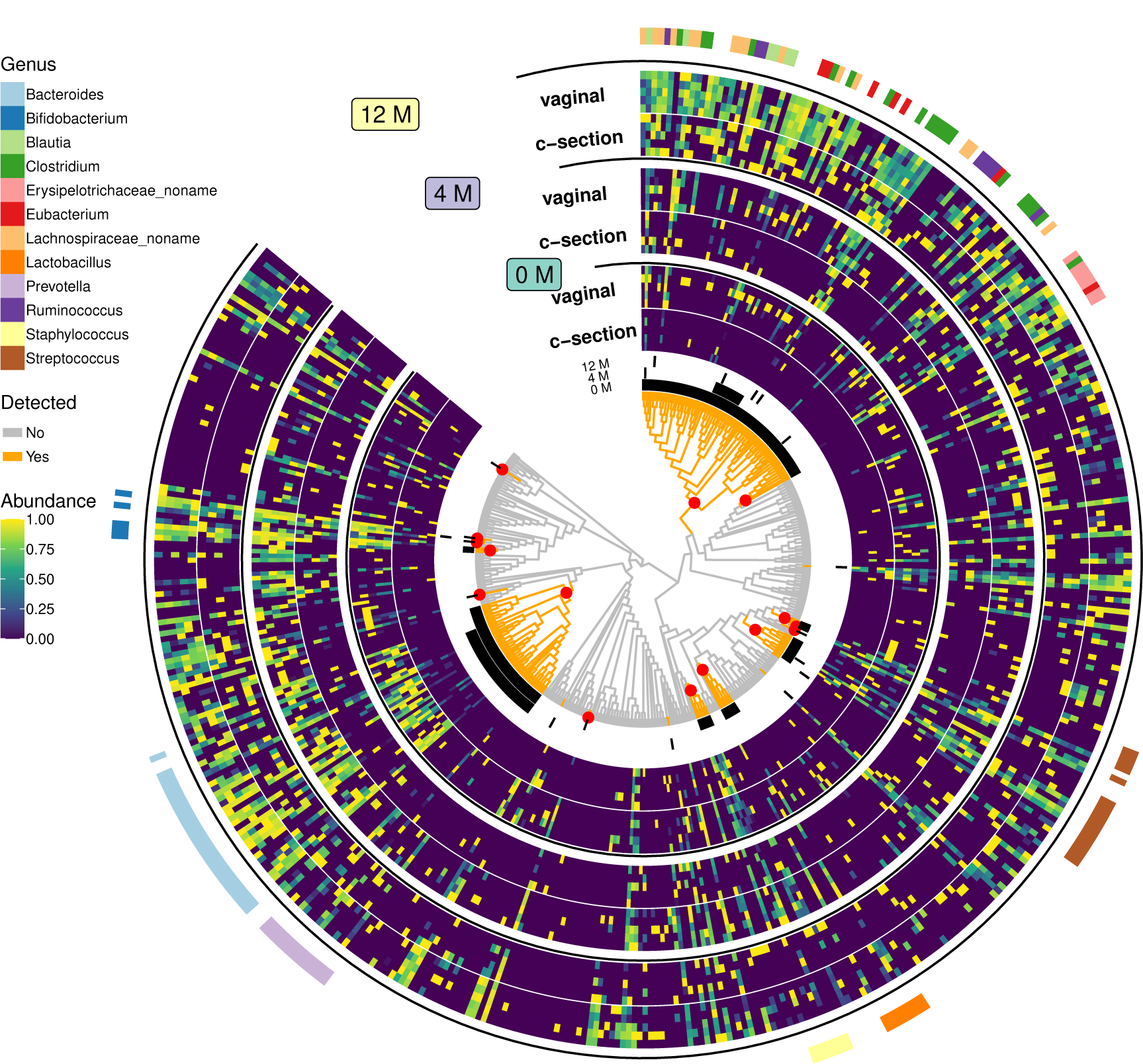
The differences of gut microbiotas between babies born vaginally and those by C-section. A phylogenetic tree of 464 metaOTUs is shown in the innermost circle. MetaOTU abundance collected from newborns at three time points: 4 days (0M), 4 months (4M), and 12 months (12M), are shown in the heatmap with log-abundance normalized to [0,1] over samples. To avoid displaying all (381) samples, average samples that are generated by randomly assigning samples within a group into 5 categories and averaging counts of each metaOTU within each category, are used. Branches detected to be DA at OM, 4M and 12M, are shown in the circular black bars between the heatmap and the tree. On the tree, orange branches are those detected to be different at least in one time point. Red dots show nodes found by *treeclimbR* for newborns (0M). Genera that have at least 5 metaOTUs reported are shown.

#### miRNA expression analysis of cardiac pressure

Similar to microbial sequences, miRNAs can be organized in a tree structure, determined not by their similarity but by their biogenesis (Figure 5). To investigate whether miRNAs with the same origin are differentially co-expressed between mice receiving transaortic constriction (TAC) or mice receiving sham surgery (Sham), we ran *treeclimbR* on a subset of the dataset from Kokkonen-Simon *et al.* [27] (see Methods). Comparison of miRNA expression between the two groups at 5% FDR identified 166 DA nodes, representing 1250 sequences belonging to 129 miRNAs. DA nodes are identified on different levels of the hierarchy: 8 genomic clusters, 16 primary transcripts, 19 miRNAs, and 123 sequences. DA branches with at least 10 descendant leaves are annotated. Those labeled with *mixed* include miRNAs of different families, which are nonetheless transcribed from genomically-clustered loci (see Supplementary Table 1). While many of the identified miRNAs had previously been reported in relation to cardiovascular health and disease [28, 29, 30, 31, 32], our analysis highlights that most of the alterations in miRNA abundance is transcriptional, including the transcriptional coregulation of genomic clusters containing mixed miRNA families, suggesting a common reshaping of chromatin at these regions.

**Figure 5:**
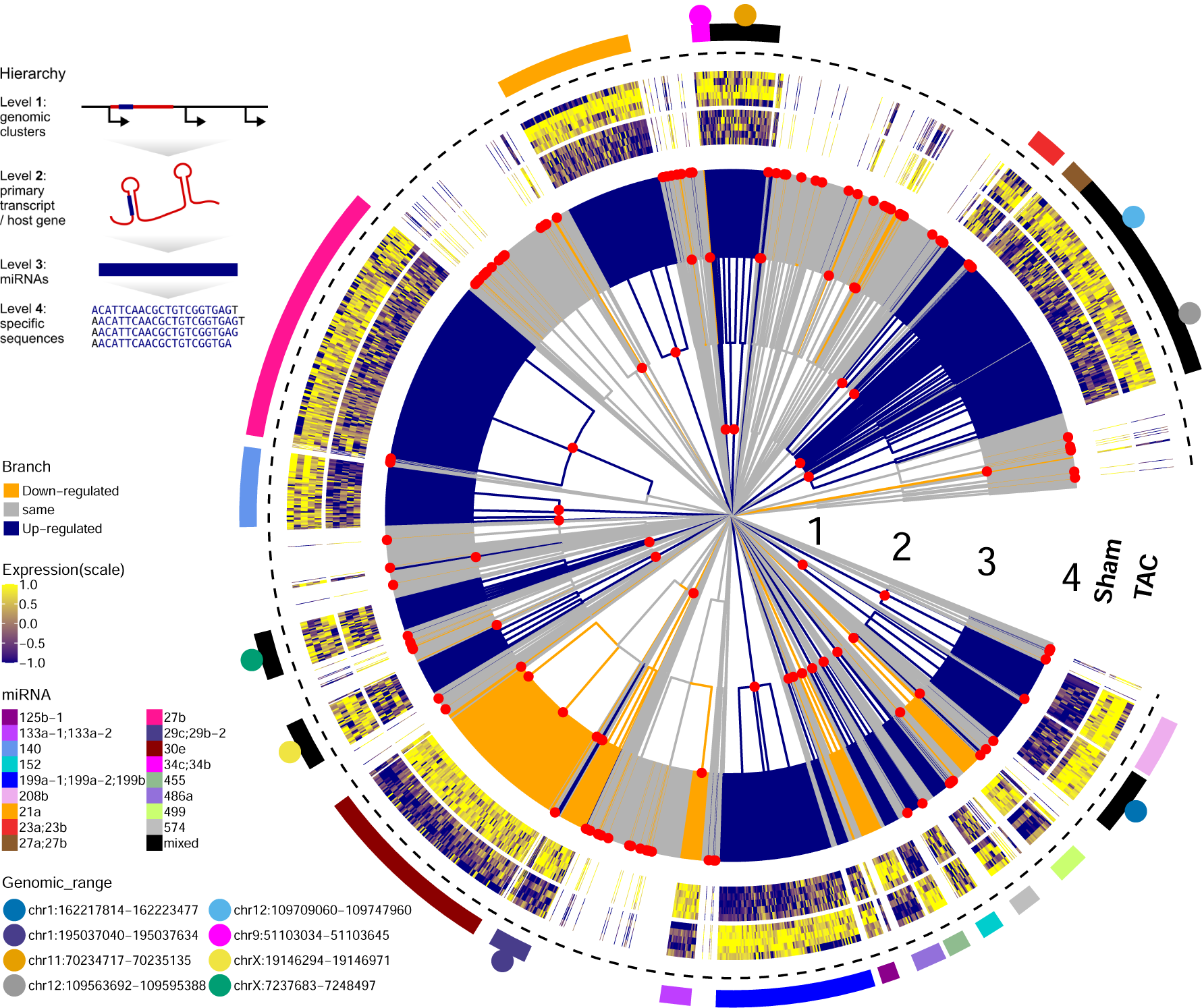
Differential expression of micro-RNAs between mice treated by sham surgery (Sham) and mice treated by transverse aortic constriction (TAC). The tree organizing the miRNAs by their origin and biogenesis is shown in the innermost circle with four levels of hierarchy: level 1 includes genomic clusters (i.e. groups of miRNAs which are located in “relatively close” regions on the genome (*<* 10*kb*)); level 2 has the primary transcript or host gene; level 3 has single-strand miRNA (e.g., mmu-miR-22-3p); level 4 has mature sequences. Nodes identified by *treeclimbR* are in red points. Up- and downregulated DA branches are shown in blue and orange, respectively. Log-expressions of sequences in identified branches are scaled and shown in the heatmap surrounding the tree. Identified branches that have at least 10 leaves are annotated, and those with more than one miRNA are labeled as mixed. The genomic ranges of identified clusters are indicated in points with different colors.

#### DS analysis of mouse cortex scRNAseq data

To explore cell state changes (DS) on a hierarchy of cell subpopulations in scRNAseq data, we applied *treeclimbR* to understand how peripheral lipopolysaccharide (LPS) affects brain cortex using 4 mice each from control (vehicle) and LPS-treated groups (see Methods). The tree that encodes the hierarchical information about subpopulations, from over-clustering, comprises 66 leaves, as shown in Figure 6a; annotation of cell types including astrocytes, endothelial cells, microglia, oligodendrocyte progenitor cells (OPC), choroid plexus ependymal (CPE) cells, oligodendrocytes, excitatory neurons, and inhibitory neurons, is taken from Crowell *et al.* [8]. Leaves within the same cell subpopulation share similar patterns according to so-called “type” markers, and mostly appear in the same branch.

**Figure 6:**
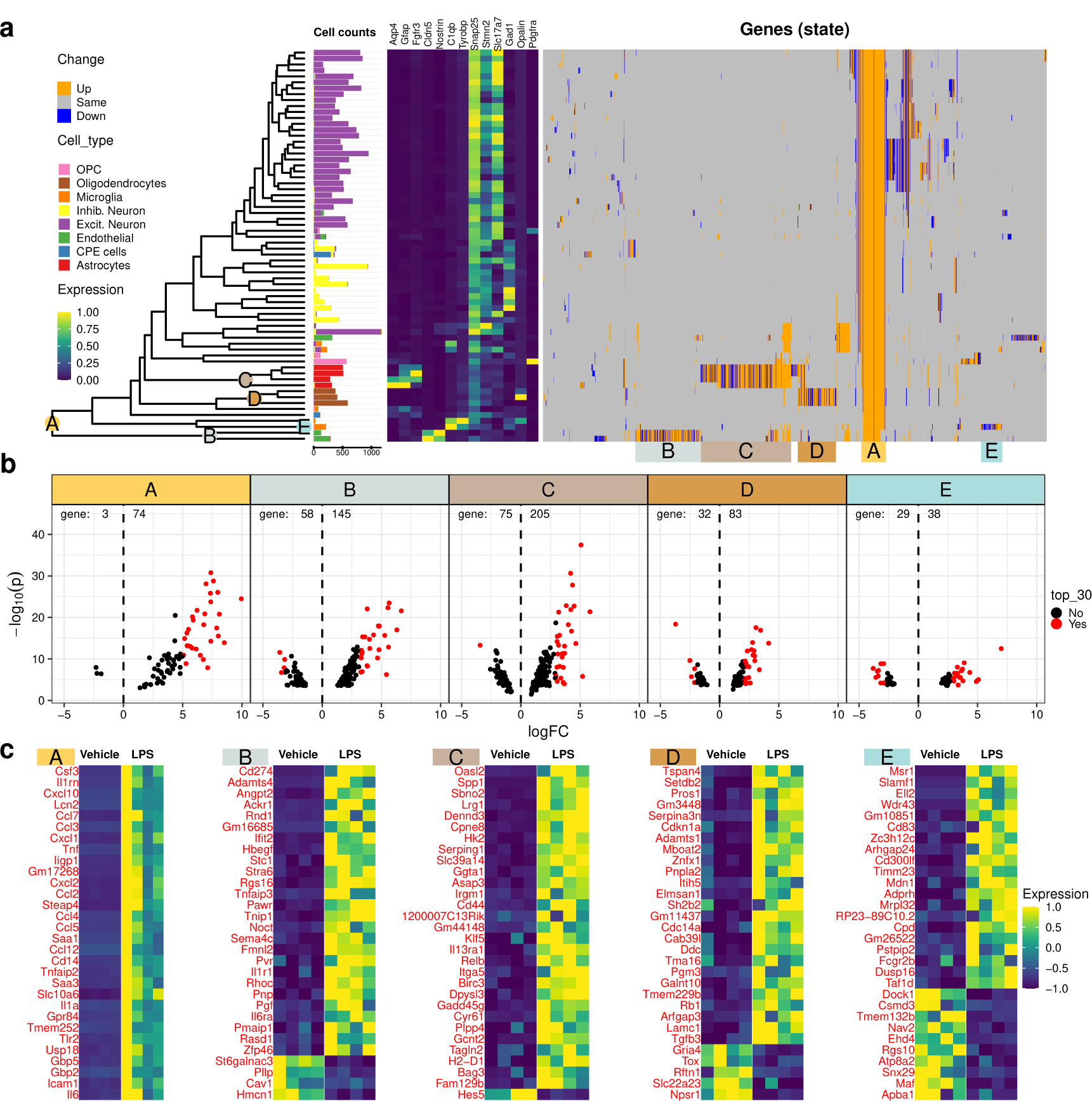
Results of comparing cortex tissue from vehicle and LPS-treated mice at different resolution levels of cell types. **(a)**. A tree encodes information of cell subpopulations at different resolutions. The cell numbers for each subpopulation are given for each leaf. The median expression of 13 canonical type markers, each columnwise scaled to [0, 1], is shown in the left heatmap. The identified DS genes are shown in the right heatmap (each column is a gene, each row subpopulation; up-regulated in LPS group in orange, down-regulated in blue). **(b)**. Volcano plots (-log P-value versus log-fold-change) with the 30 genes with highest absolute logFC colored in red. Number of down and up-regulated genes are labelled on the left and right side of the dashed line at *logFC* = 0. **(c)**. Heatmaps of top 30 genes (red points in panel **(b)**), ordered by decreasing logFC.

Using a 5% FDR threshold, *treeclimbR* identified 1561 DS genes that are expressed differently between vehicle- and LPS-treated mice in at least one cell subpopulation (genes can be deemed DS in multiple subpopulations) with absolute logFC above 1. We clustered them according to their subpopulation-wise DS pattern and summarized five distinct categories (*A − E*), as shown in Figure 6. Genes in category A change across all cell types, whereas the other four categories identify genes that change in one or two specific cell types. Since *treeclimbR* is run gene-wise (one DS test for each gene at each subpopulation), different levels of the tree can be selected for each gene. To simplify the visualization, we only label nodes where more than 70% of genes in a category were selected at the shown level. For example, the level for category *B* is selected by 99.5% of genes. Volcano plots of genes in each category, and sets with highest absolute log fold change are shown in Figure 6b and c, respectively. Inflammatory signaling has been shown to trigger the up-regulation of several cytokines in astrocytes [33], and indeed we observe the upregulation of a number of them, including Cxcl2, Cxcl1 and Ccl5, not only in astrocytes but across all cell types (category *A*).

## Discussion

Many applications in biology portray entities in a hierarchical structure. The question is then how to best leverage this information in downstream analyses where measurements (e.g., abundance) across multiple samples and experimental conditions are compared. We presented a novel principled approach, *treeclimbR*, which can be used to find a representative resolution, leading to increased power while maintaining error control. It compares favorably to leaf-level approaches (e.g., BH [21] and existing tree-based approaches (e.g., *StructFDR* [16]) when weak but coherent signals exist according to the tree.

To control FDR, *treeclimbR* assumes that leaves in branches without signal have directions up or down independently, which requires that the organization of entities on the tree is not directly driven by the changes between experimental conditions. In other words, it is recommended to have independent information on the tree and the data being analyzed. For example, in microbial data or miRNA data, the tree is organized according to sequence information (e.g., similarity or biogenesis) and the data is counts of those entities across samples. For single cell datasets, a tree can be constructed from clustering of cell type markers, and the analysis is done on state markers, although these may not be completely independent. When the same data is used for both the tree construction and the differential analysis, we might gain power to detect relevant entities while inflating the FDR due to “double dipping”. A typical example, in microbial data, is the correlation tree that is constructed based on the abundance profiles of taxa across samples from different experimental conditions. Such a tree tends to put entities showing the same direction in close proximity. In other words, it clusters not only entities with the same direction of signals in the same branch, but also those by chance appearing in the same direction. For the latter, *treeclimbR* has difficulty to distinguish it from weak but coherent signals, which overestimates the average size of signal branches *r* and the upper boundary of *t* (see Equation 6) that would further lead to poor FDR control.

Notably, the *treeclimbR* approach is flexible and users can specify any relevant method to perform the differential testing (DA and DS tests were the focus here, but other options are possible), and it may have applications beyond biology as long as P-values and estimated directions could be provided on all nodes of the tree. To successfully obtain a representative resolution, it is important that the direction of signal is correctly estimated by the chosen method. In single cell datasets, leaves of the tree (cell subpopulations) are usually obtained by unsupervised clustering, but the number of clusters is subjective and chosen according to a tuning parameter. Here, a balance needs to be struck between separating entities and having sufficient signal to allow methods to detect changes. In addition, users might need to preprocess the tree before running *treeclimbR*, for example, removing leaves or internal nodes that do not have sufficient data to reliably estimate directions of signals or, even separating a tree into multiple sub-trees, if entities (e.g., cell subpopulations) are sufficiently distinct. Taken together, *treeclimbR* is a sensitive and specific method that facilitates fine-grained inferences of hierarchical hypotheses via a rooted tree. The corresponding R package is available from https://github.com/fionarhuang/treeclimbR (will be submitted to Bioconductor) and code to reproduce all analyses is available (see Methods).

## Methods

### Simulation framework (microbiome data)

We simulate samples for two groups: control (*C*) and treatment (*T*), and generate OTU counts (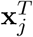 or 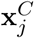) in a sample *j* from a Dirichlet-multinomial (DM) distribution with parameters estimated from a real microbial dataset, as has been suggested in several articles [16, 17]. The real throat data, *throat_v35* is subset from *V35* that is provided in the R package *HMP16SData* [34], *by* taking 153 samples collected from throat and 956 OTUs (operational taxonomic units) with non-zero count in more than 25% of samples. In particular, we sample:

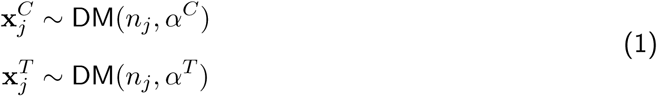

where 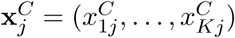and 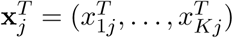 are counts of *K* = 956 OTUs in a sample *j* that belongs to control or treatment group, respectively; *n*_*j*_ is the total count of sample *j* that is randomly sampled from sequencing depths of 153 samples in *throat_v35* ; 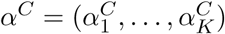and 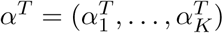 are parameters storing information about the relative abundance (proportion) and dispersion of OTUs in the control and treatment group, respectively. We estimate *α*^*C*^ using the R package *dirmult* [35] that reparameterizes *α*^*C*^ with 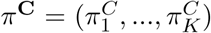 and *θ*, where 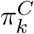 is the expected proportion of OTU *k* in a sample belonging to the control group, and *θ* is a parameter about OTU correlation. In short, 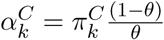 In our simulation, *θ* is estimated from *throat_v35* to apply in both control and treatment groups, and *π*^*C*^ and *π*^*T*^ are manipulated to create three scenarios: *BS, US* and *SS* (see Figure 2). The simulated data (in the control group) is shown to have similar mean-variance relationship but a bit less random zeros when compared to the real data using *countSimQC* [36] (see Supplementary countSimQC report).

In *BS*, signals are simulated on two randomly selected branches (*A* and *B*) by swapping their proportions in the treatment group as Equations 2; *US* and *SS* are in Supplementary Note 1.

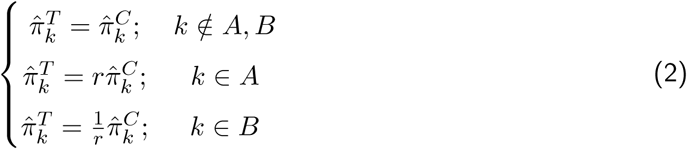

where 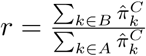 is the fold change; 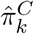 is the estimated proportion of OTU k from *throat_v35*. In other words, *π*^*C*^ is estimated from *throat_v35*, and *π*^*T*^ is obtained based on *π*^*C*^ by changing values of selected OTUs.

### Description of treeclimbR methodology

#### Data aggregation

Here, the aggregation is shown in Equation 3 and 4 for the DA and DS case, respectively. Depending on the dataset and method used in the differential analysis, mean or median might be used instead of sum. In the DA case, counts of *K* entities in *J* samples are observed and a tree on entities is constructed such that each entity can be mapped to a leaf. Data is aggregated in a way that the count of node *i* in sample *j, Y*_*ij*_ is generated as

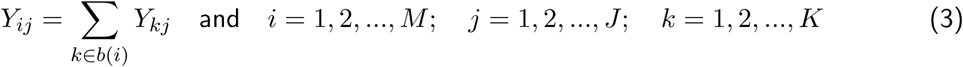

where *b*(*i*) represents the descendant leaves of node *i* (see tree notations in Figure 1); *M* is the total number of nodes on the tree; *J* is the number of samples; *K* the number of entities observed. In the DS case, we have values of *G* features observed on each cell from *J* samples, and a tree about cell subpopulations (entities) is constructed such that multiple cells are mapped to a leaf. Samples are collected from different experiment conditions. The value of feature *g* on node (cell subpopulation) *i* in sample *j*, 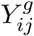, is aggregated from cells as

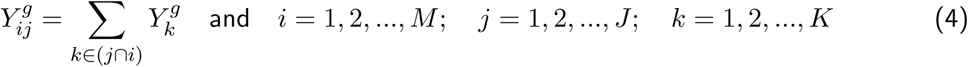

where *k* ∈ (*j* ∩ *i*) means that a cell *k* is from sample *j* and belongs to subpopulation *i* (cell *k* is mapped to the descendant leaves of node *i*, or *k* ∈ *b*(*i*)); *M, J* and *K* correspond to the total number of nodes, samples and cells, respectively.

#### The generation of candidates

Candidates are used to capture the latent signal pattern on the tree. The search for candidates is based on a *U* score defined as Equation 5:

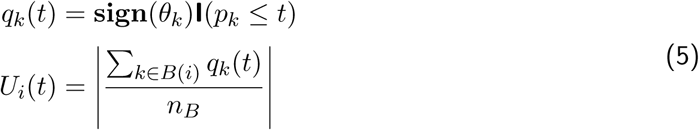

Here, *q*_*k*_(*t*) is a score of node *k*, derived from its P-value *p*_*k*_ and estimated direction **sign**(*θ*_*k*_), under a tuning parameter *t*. When *p*_*k*_ ≤ *t, q*_*k*_(*t*) = 1 with **sign**(*θ*_*k*_); otherwise, *q*_*k*_(*t*) = 0. The *U* score of node *i* at *t, U*_*i*_(*t*), is the absolute average *q* scores over nodes in *B*(*i*) that includes node *i* and its descendant nodes. *n*_*B*_ is the number of nodes in *B*(*i*). The *U* score could be considered as a measure of coordinate change within a branch. It achieves 1 when a consistent pattern, which includes both signs in the same direction and P-values below *t*, is observed; and it is close to 0 when nodes in a branch highly disagree on either the sign or P-value. With a suitable *t* value, we might expect signal branches are in a consistent pattern while others that have P-values following a uniform distribution [0, 1] and directions arbitrary up or down on leaves, are not. Since signal branches are unknown in reality, we cannot directly determine the value of *t*. To suggest different candidates of signal branches, the tree is explored by tuning *t* in the range [0, 1].

A candidate at *t* is obtained using the procedure below:

1. It starts from the root and moves toward leaves along edges.
2. For each path, it stops when a node *i* having *U*_*i*_(*t*) = 1 and *p*_*i*_ *<* 0.05 appears or the leaf is reached.

If a branch without signal by chance has the same direction, its branch node might reach *U* = 1 at high *t* (e.g., *t* = 1). In branches without signals, to keep candidate close to the leaf level, we hinder the selection of an internal node with a restriction *p*_*i*_ < 0.05. This means the probability of representing a three-leaf branch, without signals, using an internal node is around 0.01, and is much lower for a larger branch.

If multiple features exist, the procedure is carried out separately for each feature, and the overall candidate at *t*, **C**(*t*), is defined as

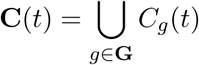

where *C*_*g*_(*t*) is the candidate of feature *g* generated at *t*, and *G* includes all features.

#### The selection of candidates

Correction for multiple testing is performed separately on each candidate, but FDR is controlled on the leaf level by limiting *t* in the range as below (see Supplementary Note 2).

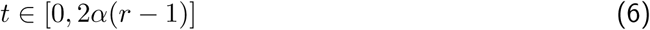

where *α* is the nominal FDR; *r* is the average size of signal branches identified at FDR = *α*. The branch size is the number of leaves in a branch. If *r* = 1, signals do not cluster on the tree, and the leaf level (*t* = 0) should be used. In real data, *r* is unknown and is estimated for a candidate *C*(*t*) as

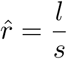

where *s* is the number of nodes with *H*_0_ rejected on the candidate **C**(**t**), and *l* is the number of descendant leaves of those rejected nodes.

Candidates that are generated with 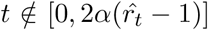 are firstly discarded to control FDR; Those that have reported the highest number of leaves with the lowest number of nodes are then selected to increase power while keeping results as short as possible.

### The preprocessing and analysis of datasets

#### Available methods

For *LEfSe*, the default settings of *LEfSe* that is installed with *conda* in *python 2.7* are used. For *miLineage*, we have applied both one-part (*miLineage1*) and two-part analysis (*miLineage2*) using the R package *miLineage v2.1*. For *lasso*, we build lasso-regularized logistic regression models, which consider values of features (e.g., abundance or expression) on all nodes of the tree as the explanatory variables, and the sample information (e.g, control or treatment group) as the response variable, with R package *glmnet 2.0-18* and chose model that gives the minimum mean cross-validated error. For *diffcyt* (*v1.6.0*), we use *diffcyt*’s *testDA_edgeR* and *testDS_limma* to analyze *AML-sim* and *BCR-XL-sim* datasets, respectively. For *StructFDR* and *HFDR*, R packages *StructFDR v1.3* and *structSSI v1.1.1* are used. Inputs on nodes (e.g., P-values) required by methods *StructFDR, HFDR, treeclimbR* and *minP* (see Supplementary Note are estimated by *edgeR v3.28.0* (*treeclimbR* ‘s *runDA* function) in all datasets, except that *diffcyt’s testDS_limma* was used in BCR-XL-sim datasets. Unless specified, the default settings provided in R packages are used for all methods.

#### Parametric synthetic microbial data

Datasets are simulated using the R package *treeclimbR* ‘s *simData* function with two randomly selected signal branches. Due to the swap of relative abundances between branches, the absolute logFC in *BS, SS* and *US* are 1.45, 2.26 and in the range [0.02, 2.13], respectively. For each scenario, five repetitions that are on the same signal branches but different counts on OTUs are made. To perform DA analysis, data was aggregated using Equation 3.

#### AML-sim and BCR-XL-sim

Datasets were downloaded from the *HDCytoData* [37] R package. According to cell type markers, cells were first grouped into a large number of clusters (400, 900, 1600 in AML-sim datasets; and 100, 400, 900 in *BCR-XL-sim* datasets) using *Flow-SOM* [38]. Then, among clusters, pairwise euclidean distances were computed using their median expressions of type markers to generate a dissimilarity matrix. Finally, the hierarchical clustering from *stats’s hclust* [39] was applied on the matrix to create a tree on clusters.

#### Infant gut microbiota data

The data was downloaded from the *curatedMetagenomicData* [40] package that provides uniformly processed human microbiome data. Only samples from babies were used. This includes a count matrix with 464 metaOTUs in rows and 381 samples in columns, and a phylogenetic tree that has 464 leaves (metaOTUs) and 463 internal nodes. Samples belong to four time points: 4 days (0M), 4 months (4M) and 12 month (12M). At each time point, there are 15 samples from the C-section group and about 80 samples (80 in 0M, 81 in 4M and 79 in 12M) from the vaginal group. Data was aggregated according to Equation 3.

#### Mouse miRNA data

The data is from Kokkonen-Simon *et al.* [27], and 10 samples, including 5 receiving TOC and 5 receiving Sham surgery, are used. The trimming, alignment and quantification of miRNA sequences were processed using *sports* [41], which ended up with 6375 miRNA sequences with counts in more than one sample. The tree was constructed based on the origins of the miRNA sequences: the miRNAs were grouped by primary transcript using the miRBase v22.1 annotation, and primary transcripts less than 10kb apart were further grouped into genomic clusters. It has 774 internal nodes and 6375 leaves. A leaf represents a unique sequence, and an internal node represents multiple sequences that share the same biological origin on a specific level. Data was aggregated as Equation 3, and *edgeR* [42] was used to compare abundance between mice receiving TOC and mice receiving Sham surgery.

#### Mouse cortex scRNAseq data

We followed the preprocessing done by Crowell *et al.* [8] that annotates cells with 8 cell types. To obtain cell type markers, expressions of genes among cell types were first compared using *FindAllMarkers* (from *Seurat v3.1.1*) separately in each vehicle-treated sample to avoid selecting LPS-related state genes; For each cell type, the top 20 genes (ranked by absolute logFC) with absolute logFC above 0.5 were then selected; We further removed markers that were only identified in one sample and finally obtained 125 marker genes. Based on 135 unique marker genes (13 canonical type marker genes and 125 computationally-identified marker genes), a tree that encodes information of cell subpopulations at different resolutions was constructed using *Seurat’s FindClusters* (resolution at 6) and *BuildClusterTree*. The tree has 66 leaves, each of them representing a cell subpopulation. To perform DS analysis, data was aggregated as Equation 4.

#### Software specifications and code availability

All analyses were run in R v3.6.2 [39]. Results were visualized with *ggplot2* [43], *ggtree* [44] *and our R package TreeHeatmap* (https://github.com/fionarhuang/TreeHeatmap). Codes to reproduce results of this study are available at https://github.com/fionarhuang/treeclimbR_article. The aggregation of tree-structure data, and the implementation of *treeclimbR* algorithm are based on R packages *TreeSummarized-Experiment* in Bioconductor and *treeclimbR* in GitHub: https://github.com/fionarhuang/treeclimbR, respectively.

## Supporting information

Supplementary (countSimQC report)

## Acknowledgments

The authors thank Dr. Lukas M. Weber (Johns Hopkins University) for assistance in preprocessing two semi-simulated CyTOF data, and members of the Robinson Lab at the University of Zurich for valuable feedback on methodology.

## Author contributions

RH, CS and MDR developed *treeclimbR*; RH, CS, TSS, CVM and MDR developed *minP* method. RH implemented methods, the simulation framework, and the method comparison; CS assisted in several technical and conceptual aspects. RH performed data analysis and interpretation of CyTOF and microbial data; RH and PLG performed data processing, analysis and interpretation of miRNA and scRNAseq data; RH and MDR drafted the manuscript, with contributions from all authors. All authors read and approved the final manuscript.

