## Supplementary Material for "treeclimbR pinpoints the data-dependent resolution of hierarchical hypotheses"

### Supplementary Figures

#### 1.1 Results of parametric synthetic microbial datasets

As shown in Supplementary Figure 1, nodes identified by *lasso*, *miLineage* are nested, where *miLineage1* and *miLineage2* are the one-part and two-part analyses provided by *miLineage*.

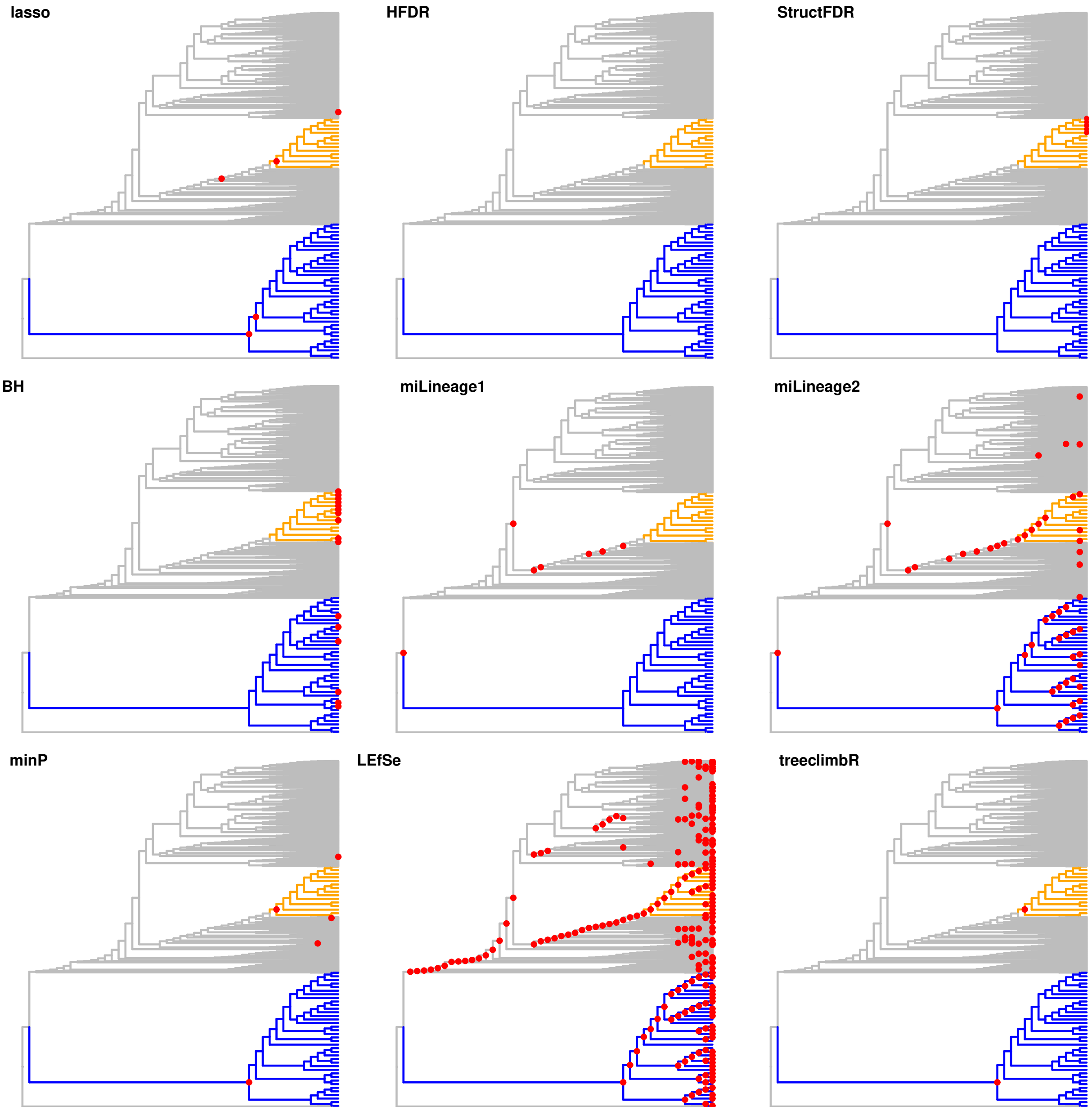

**Supplementary Figure 1. Results of methods in BS with 25 samples per group.** Two signal branches are colored in blue and orange. Nodes identified by methods are labeled as red points in each panel. *LEfSe*, *lasso* and *miLineage* (*miLineage1* and *miLineage2*) identify nodes that have ancestor-descendant relationship.

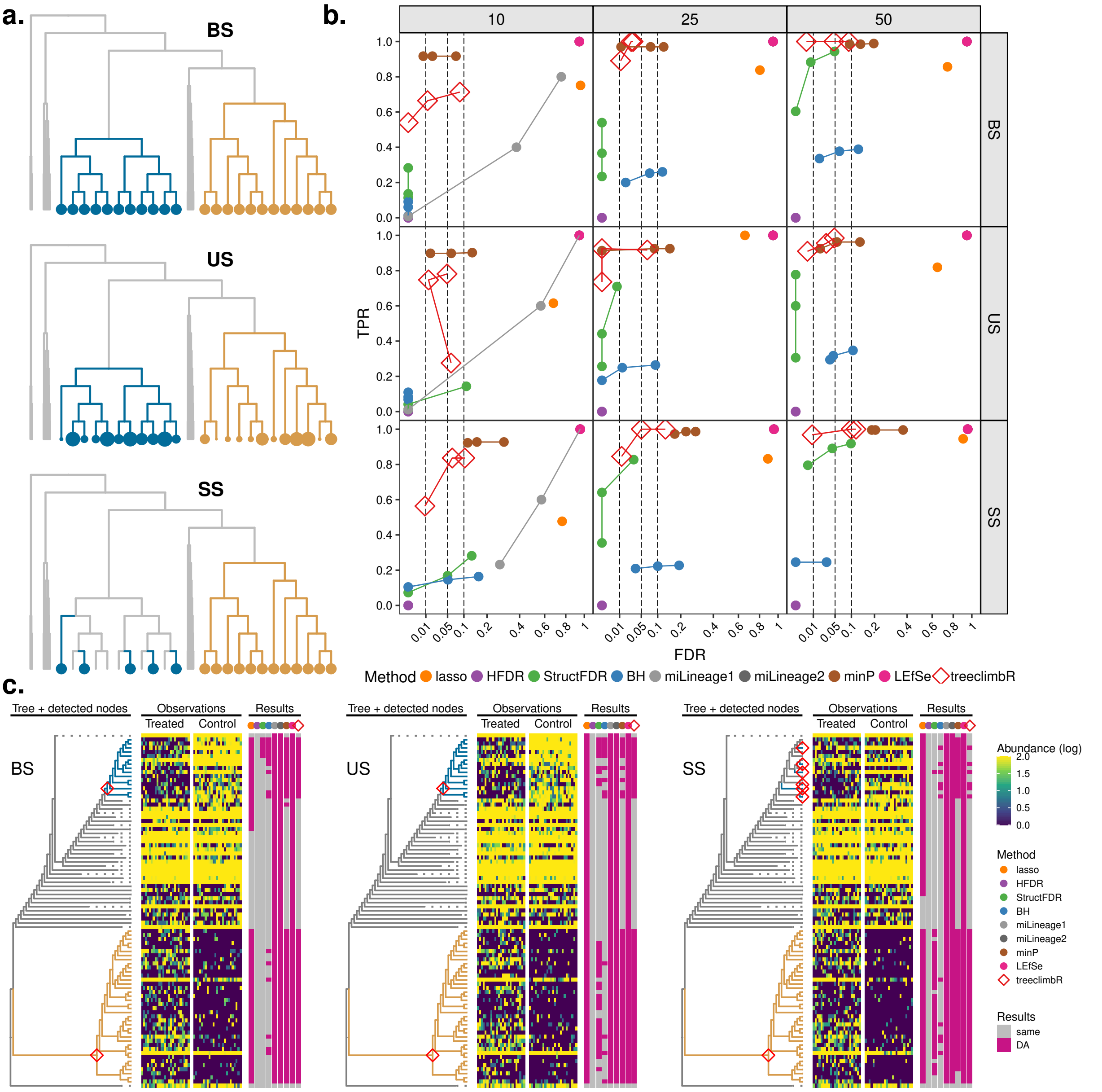

**Supplementary Figure 2. The performance of methods on parametric simulated microbial data. (a)**. A schematic example of three simulated scenarios: *BS*, *US* and *SS*. Signal branches (i.e., with DA between groups) are in turquoise (decreased) and gold (increased), and larger points represent bigger change. **(b)**. Average TPR and FDR (over 5 repetitions) of methods on three scenarios under different sample sizes (10, 25, and 50 per group). Methods are in colors. Each method has three points that represent imposed FDR cutoffs at 0.01, 0.05 and 0.1. **(c)**. DA branches identified by methods in one of 5 repetitions. The non-DA branches of the phylogenetic tree are represented with dashed lines to save space. Nodes identified by *treeclimbR* are labeled on the tree; and other methods are in Supplementary Figure 1. OTU counts are in heatmap with samples split by groups. All OTUs (rows of heatmap) identified by methods are summarized in the results panel.

Supplementary Figure 2 is similar to Figure 2, except that results of methods (*LEfSe*, *lasso* and *miLineage*) providing nested nodes are interpreted on broad resolution levels. Figure 2 uses identified nodes that have no descendant nodes identified; Supplementary Figure 1 uses identified nodes with their ancestors not identified. For example, in the *lasso* panel of Supplementary Figure 1, two nodes (a parent node and a child node) are identified in the blue branch, the child is used in Figure 2 and the parent is used in Supplementary Figure 2. Therefore, in Supplementary Figure 2*b*, *lasso* has higher TPR but also much higher FDR than as shown in Figure 2. For *miLineage*, it has TPR equal to 1, but also FDR equal to 1 in most simulations. In other words, it overtakes signal branches and most identified branches are false positives. This could also be seen in the *miLineage1* and *miLineage2* panels of Supplementary Figure 1.

#### 1.2 Results of non-parametric synthetic microbial datasets

Bichat *et al.* [1] have performed non-parametric simulations based on a real gut dataset from Brito *et al.* [2]. They first assigned 112 samples randomly to two groups, selected $m$ taxa randomly from the most prevalent taxa, and multiplied counts of the $m$ taxa in one of the two groups by a fold change $fc$. In other words, the tree is uninformative in the simulation. Different values of $fc$ and $m$ have been implemented and results are shown in Figure 5 of Bichat *et al.* [1], highlighting that a tree-based method, *StructFDR*, has no gain and even performs worse (lower power) than the classical procedure *BH* when using the non-informative tree (taxonomic tree). To test whether our tree-based algorithm *treeclimbR* has the same issue, we reproduced their work and added *treeclimbR* to the comparison using the taxonomic tree. Results are in Supplementary Figure 3, which shows that *treeclimbR* performs similarly to *BH* when the tree is uninformative.

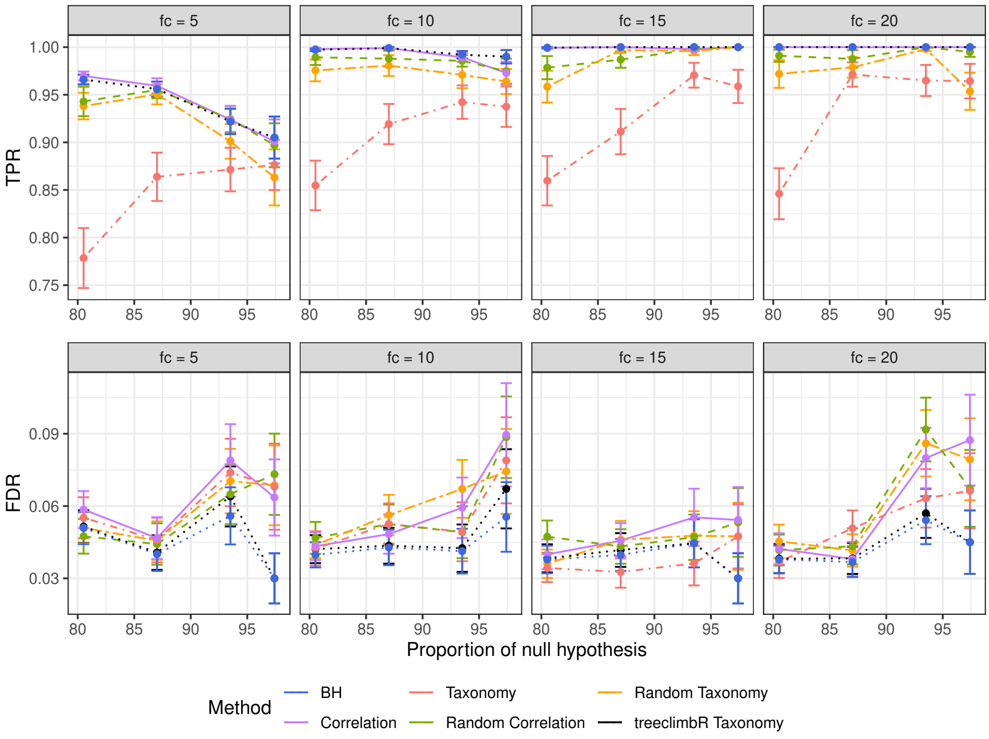

**Supplementary Figure 3. Results of different FDR control procedures on non-parametric simulation.** Mean and Squared Error of the Mean (SEM) of the true positive rates (TPR, top) and FDR (bottom) per different fold changes (facets) for non-parametric simulations. The different FDR control procedures are color-coded. Mean and SEM are computed over 100 replicates.

#### 1.3 Results of BCR-XL-sim datasets

In addition to *pS6*, protein markers including *pPIcg2*, *pErk*, *pNFkB* are also differentially expressed in B cells when comparing stimulated samples against control samples. In Supplementary Figure 4, we see that only *treeclimbR* identifies the big branch of B cells for three markers; other methods either find some sub-branches or miss the whole big branch of B cells. By scanning the whole tree to find the suitable resolution, *treeclimbR* could perform well on high-resolution clusters (e.g. 2500 clusters) that becomes difficult for *diffcyt* (See Supplementary Figure 4d).

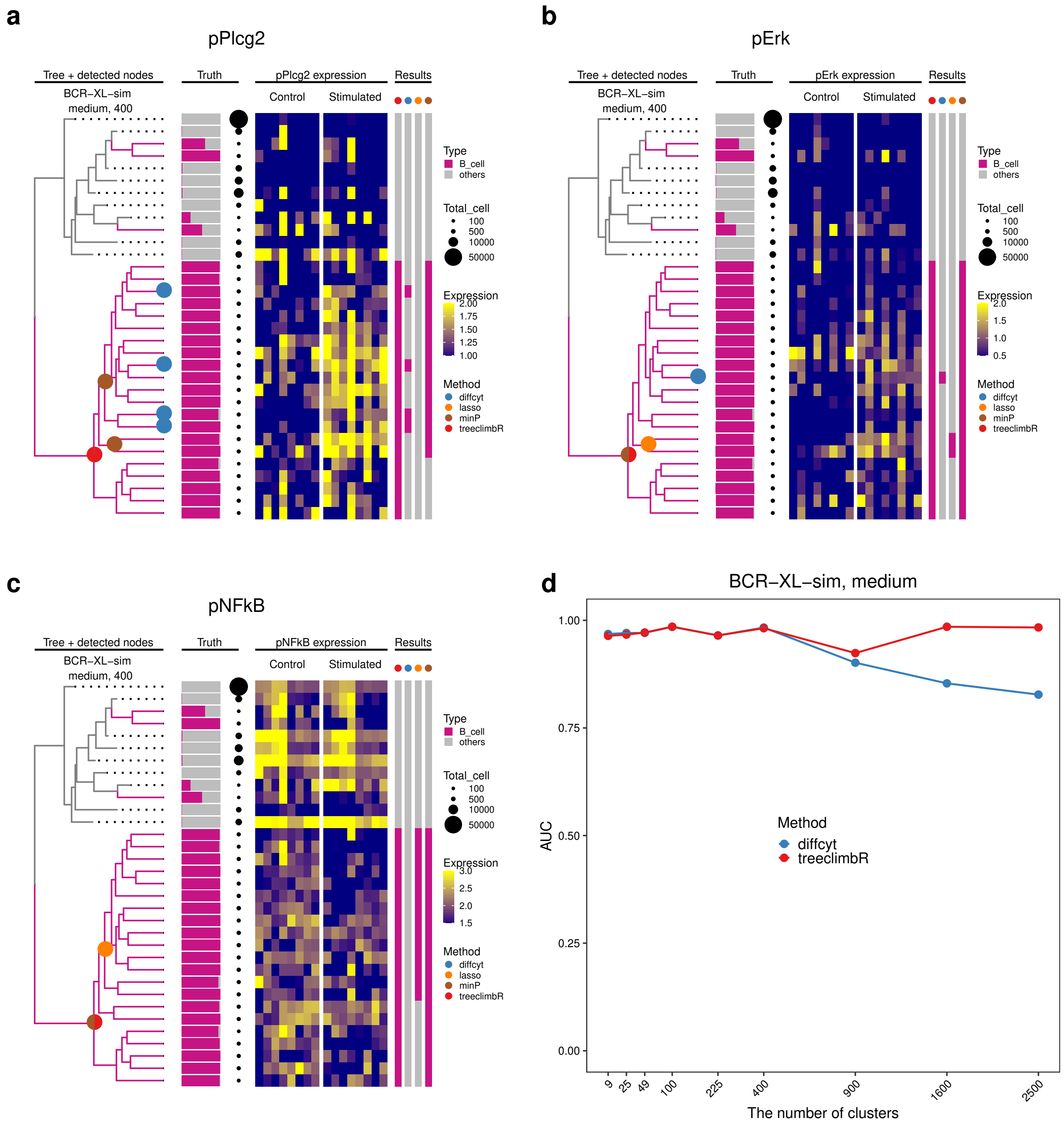

**Supplementary Figure 4. Results of methods on the medium scenario of *BCR-XL-sim*.** Branches that are identified to have differential expression of *pPlcg2*, *pErk* and *pNFkB* in the medium scenario of *BCR-XL-sim* are shown correspondingly in plots **a, b, c**. Each plot has four panels showing tree, truth, observations and results. Non-DS branches (without B cells) are represented in dashed lines to save space; DS branches (with B cells above 50%) are in purple. Nodes identified by methods are in different colors. The true cell-type compositions and cell counts (point sizes) on each leaf are given in the truth panel. The median expressions of marker on leaves are shown as heatmap with cell subpopulations in rows and samples in columns (split by groups). Descendant leaves of identified nodes are annotated in the results panel. **d** Area under ROC curves (AUC) of *diffcyt* and *treeclimbR* when using different numbers of clusters (between 9 and 2500 clusters).

#### 1.4 Results of miRNA data

| Region | miRNAs |
|---|---|
| chr1:195037040-195037634 | miRNA-29C, miRNA-29b-2 |
| chr1:162217814-162223477 | miRNA-199a-2, miRNA-214 |
| chr11:70234717-70235135 | miRNA-195a, miRNA-497a |
| chr12:109563692-109595388 | miRNA-127, miRNA-136, miRNA-337, miRNA-431, miRNA-433, miRNA-434, miRNA-540, miRNA-673 |
| chr12:109709060-109747960 | miRNA-134, miRNA-154, miRNA-299a, miRNA-300, miRNA-329, miRNA-369, miRNA-376, miRNA-377, miRNA-379, miRNA-380, miRNA-381, miRNA-382, miRNA-409, miRNA-410, miRNA-411, miRNA-485, miRNA-541, miRNA-667, miRNA-668 |
| chr9:51103034-51103645 | miRNA-34c, miRNA-34b |
| chrX:19146294-19146971 | miRNA-221, miRNA-222 |
| chrX:7237683-7248497 | miRNA-188, miRNA-362, miRNA-500, miRNA-501, miRNA-532 |

**Supplementary Table 1. miRNAs in 8 identified genomic clusters.**

### Supplementary Note 1: Simulation framework

In Supplementary Figure 5, the height of the bar represents the proportion of entities (in the specified branch) within a sample. We consider the tree structure as three parts, the two selected DA-branches and the rest. The phenotypic outcome is represented by the group (control or treatment). The proportions of the orange and blue branches are swapped between different groups in scenario *BS* and *US*. The height of the orange bar in the treatment group is equal to the blue bar in the control group, and the height of the blue bar in the treatment group is the same as that of the orange bar in the control group. In *BS*, the increase or decrease in the proportion is evenly distributed to the entities within branch by a common factor $r$ (in the blue branch) or $\frac{1}{r}$ (in the orange branch). However, in *US*, the change of proportion is not evenly distributed and the factor $r_k$ varies on entities. In *SS*, the swap is partial and depends on the value $c$. That's why the height of the orange bar in the treatment group is different to that of the blue bar in the control group.

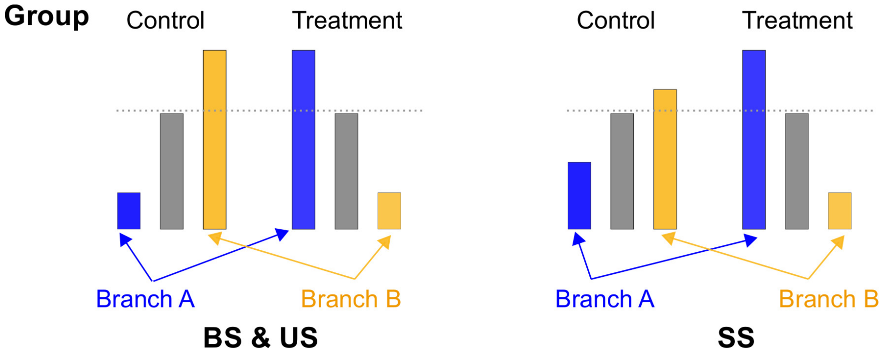

**Supplementary Figure 5. The overall idea of simulated scenarios.**

#### 2.1 Balanced signal: BS

**Proof:** The proportions of two branches are swapped between groups in BS.

$$\begin{aligned}
\sum_{g\in A} \hat{\pi}_g^T &= \sum_{g\in A} r_g \hat{\pi}_g^C = \sum_{g\in A} \hat{\pi}_g^C r_g \\
&= \sum_{g\in A} \hat{\pi}_g^C (1 + \frac{\sum_{k\in B} \hat{\pi}_k^C - \sum_{g\in A} \hat{\pi}_g^C}{\sum_{g\in A} \hat{\pi}_g^C u_g} u_g) \\
&= \sum_{g\in A} \hat{\pi}_g^C + \sum_{g\in A} \hat{\pi}_g^C \frac{\sum_{k\in B} \hat{\pi}_k^C - \sum_{g\in A} \hat{\pi}_g^C}{\sum_{g\in A} \hat{\pi}_g^C u_g} u_g \\
&= \sum_{g\in A} \hat{\pi}_g^C + (\sum_{k\in B} \hat{\pi}_k^C - \sum_{g\in A} \hat{\pi}_g^C) \frac{\sum_{g\in A} \hat{\pi}_g^C u_g}{\sum_{g\in A} \hat{\pi}_g^C u_g} \\
&= \sum_{k\in B} \hat{\pi}_k^C
\end{aligned} \tag{1}$$

Similarly, we could have $\sum_{k\in B} \hat{\pi}_k^T = \sum_{g\in A} \hat{\pi}_g^C$.

#### 2.2 Unbalanced signal: US

**Unbalanced signal (US)**. Two branches ($A$ and $B$)are randomly selected to swap their proportion.

$$\begin{cases} \hat{\pi}_k^T = \hat{\pi}_k^C; & k \notin A, B \\ \hat{\pi}_k^T = r_k \hat{\pi}_k^C, \quad r_k = 1 + \frac{\sum_{k \in B} \hat{\pi}_k^C - \sum_{k \in A} \hat{\pi}_k^C}{\sum_{k \in A} \hat{\pi}_k^C u_k} u_k; & k \in A \\ \hat{\pi}_k^T = r_k \hat{\pi}_k^C, \quad r_k = 1 + \frac{\sum_{k \in A} \hat{\pi}_k^C - \sum_{k \in B} \hat{\pi}_k^C}{\sum_{k \in B} \hat{\pi}_k^C u_k} u_k & k \in B \end{cases} \tag{2}$$

where $u_k$ is randomly generated for entity $k$ from a uniformly distributed function as Equation 3 and $t$ is set to make sure $r_k$ is above zero.

$$\begin{cases} u_k \sim \mathrm{U}(b, 1), \quad b = \max(1 - \frac{\sum_{k \in B} \hat{\pi}_k^C}{\sum_{k \in A} \hat{\pi}_k^C}, 0); & k \in A \\ u_k \sim \mathrm{U}(b, 1), \quad b = \max(1 - \frac{\sum_{k \in A} \hat{\pi}_k^C}{\sum_{k \in B} \hat{\pi}_k^C}, 0); & k \in B \end{cases} \tag{3}$$

The proof about the proportions of two branches being swapped is similar to Equation 1.

#### 2.3 Sporadic signal: SS

**Sporadic signal (SS)**. Two branches ($A$ and $B$)are randomly selected, but only some leaves in branch $A$ (denoted as $A'$) are randomly selected to swap their proportion with leaves in branch $B$.

$$\begin{cases} \hat{\pi}_k^T = \hat{\pi}_k^C; & k \notin A', B \\ \hat{\pi}_k^T = r \hat{\pi}_k^C; & k \in A' \\ \hat{\pi}_k^T = c \hat{\pi}_k^C; & k \in B \end{cases} \tag{4}$$

where $r = \frac{(1-c) \sum_{k \in B} \hat{\pi}_k + \sum_{k \in A'} \hat{\pi}_k}{\sum_{k \in A'} \hat{\pi}_k}$ and $0 < c < 1$. The proof about the proportions of two branches being swapped is similar to Equation 1.

### Supplementary Note 2: Details about *treeclimbR*

#### 3.1 The leaf FDR and candidates

Our algorithm *treeclimbR* performs multiple hypothesis tests on the candidate level but controls FDR on the leaf level.

**The definition of the leaf FDR.** When a null hypothesis on an internal node is rejected, null hypotheses on all its descendant leaves are considered as being rejected too. We define the leaf FDR as the ratio of the number of falsely rejected leaves to the number of rejected leaves. This is formulated as

$$\mathbf{FDR}_L = \mathbf{E}[Q_L] = \mathbf{E}[\frac{V_L}{V_L + S_L}] \tag{5}$$

where $V_L$ and $S_L$ represent the numbers of true positive and false positive leaves, respectively. A true positive leaf is a leaf with non-true null hypothesis and correctly declared as non-true null hypothesis; a false positive leaf is a leaf with true null hypothesis but wrongly declared as non-true null hypothesis;

**Examples of candidates.** In *treeclimbR*, multiple candidates are obtained by tuning a parameter $t$ to capture the signal pattern on a tree. A toy example with three candidates is shown in Supplementary Figure 6. Without using a tree, there are $K$ hypothesis tests on $K$ entities (leaves $1 \sim K$). When a tree is available, *treeclimbR* tends to represent branches that have coherent change with internal nodes. This leads to less hypothesis tests. An example is candidate 2 that replaces leaves $5 \sim 9$ with an internal node $X$. When $t$ is set too high, candidate 3 that replaces leaves $5 \sim 10$ with an internal node $Y$ might be obtained. We perform multiplicity correction on the candidate level, but aim to control FDR on the leaf level (Equation 5).

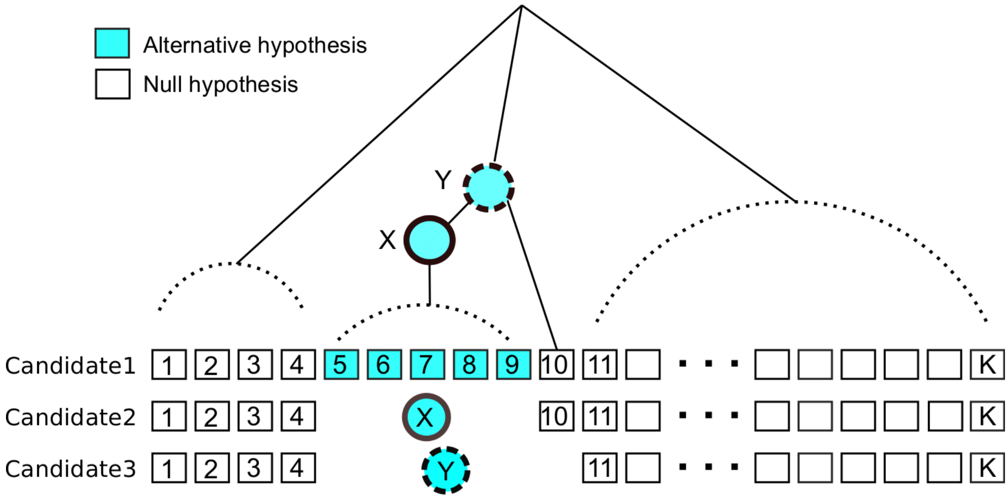

**Supplementary Figure 6. A toy example of three different candidates on the tree.** Dashed lines are used to represent branches with different possible structures. Each leaf represents an entity. Entities with true alternative hypothesis are colored in blue; otherwise in white. Node $Y$ is considered as true $H_1$ in candidate 3 because it includes leaves with true $H_1$ and its P-value wouldn't follow uniform distribution anymore. The rejection of null hypothesis on node $Y$ would contribute four true positives and one false positive in calculating the leaf FDR. Note: Directions of blue leaves are the same.

The use of candidates might improve or deteriorate the control of the leaf FDR. We know that FDR is defined as Equation 6 in the BH procedure [3].

$$\mathbf{E}[Q] = \mathbf{E}[\frac{V}{V+S}] \tag{6}$$

where $V$ is the number of false positives (true null hypotheses but declared as non-true null hypotheses) and $S$ is the number of true positives (non-true null hypotheses and correctly declared as non-true null hypotheses). When the power to detect signal leaves is the same for both candidates, $S$ of candidate 2 is lower than that of candidate 1 due to the use of internal nodes. Hence, when the FDR is well controlled in both candidates, $V$ in candidate 2 tends to be lower than candidate 1, which implies the leaf FDR is improved in candidate 2. However, it doesn't mean that candidates using internal nodes to represent signal entities always have lower FDR than the leaf candidate. The generation of candidates might introduce new false positives. For example, if null hypothesis on node $Y$ is rejected, the leaf 10 is falsely reported as positive. Such a false positive is caused by the generation of the candidate instead of the BH procedure, and is more likely to occur on candidates generated with high $t$. To address the issue, the range of $t$ needs to be limited to control the leaf FDR.

#### 3.2 The range of $t$

A tree might have multiple signal branches, each of which has different surroundings (e.g. the size of sibling branch). To simplify, let's consider one signal branch at a time. In Supplementary Figure 6, the node $Y$ is selected to represent leaves $5 \sim 10$ if all its descendants have P-values below the specified threshold $t$, and the leaf 10 occurs to have the same estimated direction as leaves $5:9$. Generally, the two descendant branches of node $Y$ could be in any configuration (e.g., more leaves or nodes are connected in different ways), and Supplementary Figure 6 shows one possible structure.

Let's say the two children of $Y$ are node $X$ and $Z$. We use $\tilde{\mathbf{H}}$ to represent the fact that all descendant leaves of $X$ are true $H_1$ and all descendant leaves of $Z$ are true $H_0$ in Equation 7. Given $\tilde{\mathbf{H}}$, the probability ($\pi$) to reject null hypothesis on node $Y$ ($H_Y$) in a candidate $\mathbf{C(t)}$ could be formulated according to Equation 7.

$$\begin{aligned}
\pi &= \mathbf{Pr}[\{H_Y \notin H_0\} \cap \{Y \in \mathbf{C(t)}\} | \tilde{\mathbf{H}}] && (7)\\
&\leq \mathbf{Pr}[Y \in \mathbf{C(t)} | \tilde{\mathbf{H}}] && (8)\\
&= \mathbf{Pr}[(\cup_{i \in B(Y)} p_i \leq t) | \tilde{\mathbf{H}}] \cdot P^* && (9)\\
&\leq \mathbf{Pr}[(\cup_{i \in b(Y)} p_i \leq t) | \tilde{\mathbf{H}}] \cdot P^{**} && (10)\\
&< \mathbf{Pr}[(\cup_{i \in b(Z)} p_i \leq t) | \tilde{\mathbf{H}}] \cdot P^{**} && (11)\\
&= \prod_{i \in b(Z)} 0.5 \cdot \mathbf{Pr}[p_i \leq t | \tilde{\mathbf{H}}] && (12)\\
&= (0.5t)^n
\end{aligned}$$

where $\mathbf{C(t)}$ is the candidate generated at $t$; $B(Y)$ includes node $Y$ and its descendant nodes; $b(X)$, $b(Y)$ and $b(Z)$ correspond to descendant leaves of node $X$, $Y$ and $Z$; $P^*$ and $P^{**}$ are probability that the estimated directions are the same for nodes in $B(Y)$ and $b(Y)$, respectively; $p_i$ is the P-value of node $i$; $n$ is the number of descendant leaves of node $Z$. Equation 7 is less than Equation 8 because the latter doesn't require $H_0$ on node $Y$ to be rejected. As discussed above, node $Y$ is selected when it and its descendants agree on the estimated direction and their P-values are below $t$. So, Equation 8 is equal to Equation 9. Equation 9 is less than Equation 10 because the latter imposes restriction on less nodes ($b(Y) \subseteq B(Y)$). Similarly, Equation 10 is less than Equation 11 because of $b(Z) \subset b(Y)$. As node $i \in b(Z)$ has true $H_0$, its P-value ($p_i$) follows uniform distribution ($p_i \sim U(0,1)$) and its probability to show the same estimated direction as leaves of $X$ is 0.5, which means Equation 11 is equal to Equation 12. Hypothesis tests on the leaf level are considered to be independent, so we could further simplify Equation 12 to $(0.5t)^n$, which decreases as $n$ increases at a fixed $t$. In other words, it becomes unlikely to select and further reject an internal node that represents a signal branch together with a sibling branch when the sibling branch is large. However, once this occurs, it brings more false positives.

The expected number of false positives that are introduced by the generation of candidates depends on the tree structure. A tree that has small non-signal branches next to signal branches is expected to have higher false positives than a tree that have large non-signal branches next to signal branches. Consider a tree that has $N$ branches with signal, the expected number of false positives ($\mathbf{E}[F]$) is maximized in the situation where each signal branch has a one or two-leaf sibling branch that has no signal. Following Equation 12, we have

$$\begin{aligned} \mathbf{E}[F] &= \sum_{i=1}^{N} n_i \cdot \pi_i \\ &< \sum_{i=1}^{N} n_i \cdot (0.5t)^{n_i} \\ &\leq N \cdot 0.5t \end{aligned} \tag{13}$$

where $n_i$ is the number of leaves on the sibling branch of the $ith$ signal branch.

Consider $t$ that is used to generate a candidate, where $s_0$ true positive nodes (covering $l_0$ true positive leaves) are reported under BH [3] procedure with FDR $\alpha$, as a variable. We would like to get the range of $t$, in which the leaf FDR is expected to not exceed $\alpha$. The FDR on the candidate and the leaf level could be formulated as Equation 14 and Equation 15, respectively.

$$\mathbf{E}[Q] = \mathbf{E}[\frac{V}{V+s_0}] \tag{14}$$

$$\mathbf{E}[Q_L] = \mathbf{E}[\frac{V_L}{V_L+l_0}] \tag{15}$$

Here, $V$ and $V_L$ are the number of false positives on the candidate and leaf level, respectively. With a fixed $l_0$, we could apply Jensen's inequality on Equation 15 to get its upper bound as Equation 16.

$$\begin{aligned} \mathbf{E}[Q_L] &= \mathbf{E}[1 - \frac{l_0}{V_L+l_0}] \\ &\leq 1 - \frac{l_0}{\mathbf{E}[V_L]+l_0} \end{aligned} \tag{16}$$

The number of false positives ($V_L$) on the leaf level comes from three different sources.

$$\mathbf{E}[V_L] = \mathbf{E}[V_L' + V_L'' + V_L'''] \tag{17}$$

- $V'_L$: False positives due to the BH [3] procedure performed on the candidate level.
- $V''_L$: False positives from the generation of a candidate (See candidate 3 in Supplementary Figure 6).
- $V'''_L$: False positives from a case similar to candidate 3 in Supplementary Figure 6) but the branch below node $X$ in Supplementary Figure 6 has fake signal. In other words, the consistent signal pattern on branch below $X$ is by chance.

Overall, $V'_L$ is due to the procedure of multiple testing correction, $V''_L$ is due to the generation of candidates, and $V'''_j$ is due to the combination of both.

With a fixed $s_0$, according to Jensen's inequality, the upper bound of Equation 14 is as below.

$$\begin{aligned}\mathbf{E}[Q] &= \mathbf{E}[\frac{V}{V+s_0}] \\ &= 1 - \mathbf{E}[\frac{s_0}{V+s_0}] \\ &\leq 1 - \frac{s_0}{\mathbf{E}[V]+s_0}\end{aligned}$$

We set the right side of the inequality below $\alpha$ to conservatively control FDR for a candidate.

$$1 - \frac{s_0}{\mathbf{E}[V]+s_0} \leq \alpha \Leftrightarrow \mathbf{E}[V] \leq \frac{\alpha}{1-\alpha}s_0 \tag{18}$$

So, the upper bound of false positives allowed in the multiple testing correction is $\frac{\alpha}{1-\alpha}s_0$. According to Equation 13 and Equation 17, we know that

$$\mathbf{E}[V'_L] = \mathbf{E}[V] \leq \frac{\alpha}{1-\alpha}s_0 \tag{19}$$

$$\mathbf{E}[V''_L] \leq \frac{s_0 t}{2} \tag{20}$$

$$\mathbf{E}[V'''_L] \leq \frac{\alpha}{1-\alpha}s_0\frac{t}{2} \tag{21}$$

Therefore,

$$\begin{aligned}\mathbf{E}[V_L] &= \mathbf{E}[V'_L] + \mathbf{E}[V'''_L] + \mathbf{E}[V''_L] \\ &\leq \frac{\alpha}{1-\alpha}s_0 + \frac{\alpha}{1-\alpha}s_0\frac{t}{2} + \frac{s_0 t}{2} \\ &= \frac{\alpha s_0}{1-\alpha} + \frac{s_0 t}{2(1-\alpha)}\end{aligned} \tag{22}$$

By substituting Equation 22 to Equation 16, we get

$$\mathbf{E}[Q_L] < 1 - \frac{l_0}{\frac{\alpha s_0}{1-\alpha} + \frac{s_0 t}{2(1-\alpha)} + l_0} \tag{23}$$

$$= \frac{2\alpha + t}{2\alpha + t + 2(1-\alpha)\frac{l_0}{s_0}} \tag{24}$$

To control the leaf FDR, we could conversely restrict the upper bound of $\mathbf{E}[Q_L]$ to be below $\alpha$.

$$\frac{2\alpha + t}{2\alpha + t + 2(1-\alpha)\frac{l_0}{s_0}} \leq \alpha \tag{25}$$

This is further simplified as

$$t \leq 2\alpha(\frac{l_0}{s_0} - 1)$$

To summarize, $t$ is non-negative and its range should be

$$t \in [0, 2\alpha(r-1)]$$

where $r = \frac{l_0}{s_0}$ is the average size of signal branches that are detected. If $r = 1$, signal detected does not cluster on the tree and the leaf level ($t = 0$) should be used to perform multiple test correction. In reality, $l_0$ and $s_0$ is unknown and we estimate $r$ as

$$\hat{r} = \frac{l}{s} \tag{26}$$

where $s$ is the number of rejected nodes on the candidate level and $l$ is the number of descendant leaves of rejected nodes. As false positives are less likely to cluster, $r$ tends to be underestimated using $\hat{r}$. This would lead to conservative results on the FDR control.

#### 3.3 The selection of P-values is unbiased in branches without signals

To control FDR on the candidate level when applying the BH procedure, it requires that P-values of true null hypotheses on the candidate level still follow a uniform distribution [0, 1]. In other words, the generation of candidates would not lead to a biased selection of P-values on branches without signals. To show this, we generate a negative control case with a tree that has no signal branches.

The toy data includes a random tree with 1000 leaves and a count matrix with 1000 rows and 60 columns. Each row of the matrix is linked to a leaf of the tree and represents an entity. Each column represents a sample. Counts of entities in a sample are randomly taken from a multinomial distribution as below:

$$Y_j \sim \text{Mult}(10000, p)$$

where $p = (p_1, p_2, ..., p_{1000})$ includes the proportions of entities. It is randomly generated and is the same for all samples.

To run *treeclimbR*, candidates are generated using the default values of $t$ in our *treeclimbR* package. The cumulative distribution of P-values on each candidate is shown in Supplementary Figure 7, curves of all candidates almost overlap each other, and are very close the dashed line that represent a uniform distribution in $[0, 1]$, which implies there is no bias in the selection of P-values on different candidates.

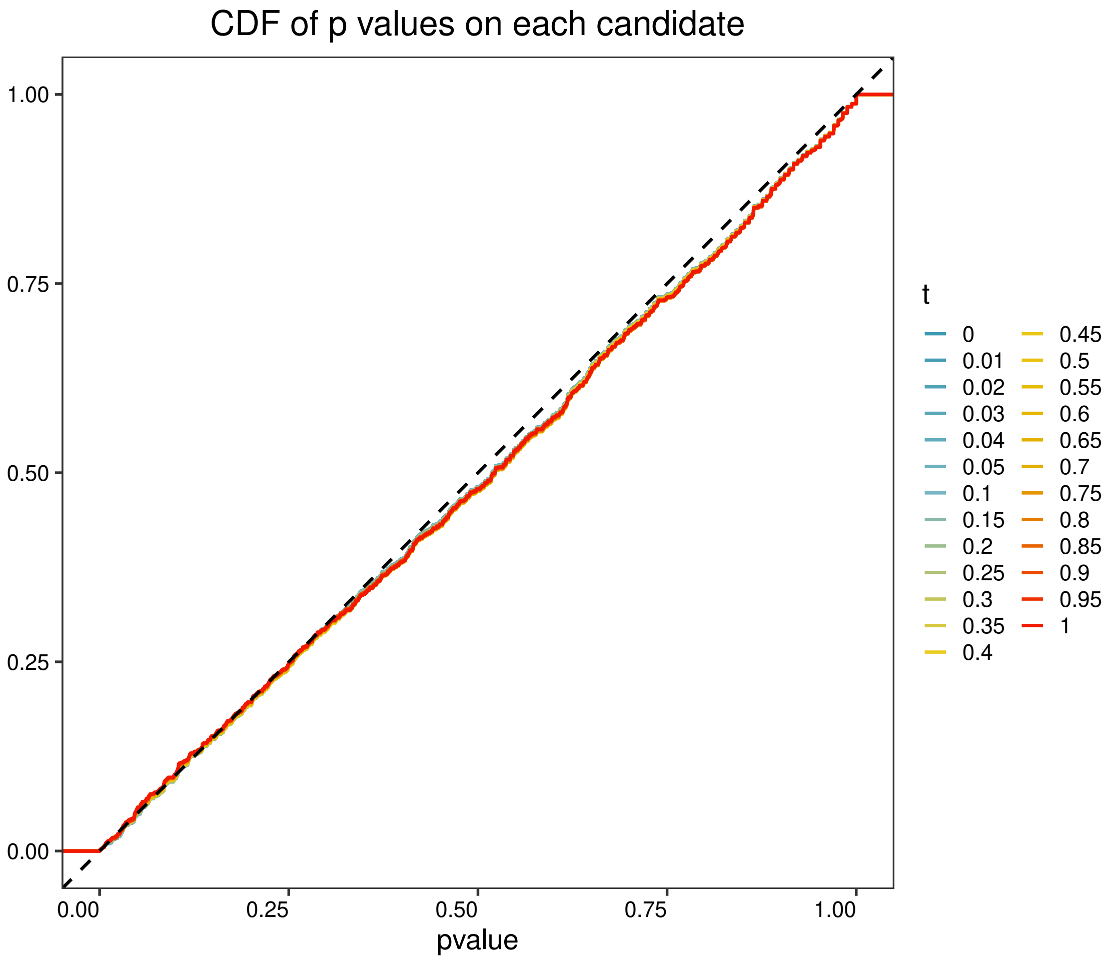

**Supplementary Figure 7. The cumulative distribution of P-values on 25 candidates.** Candidates that are generated under different $t$ values have almost the same cumulative distribution. The dashed line represent an uniform distribution in $[0, 1]$.

#### 3.4 An example on toy data

Here, we use a toy dataset to show how *treeclimbR* scans the tree under different $t$ values. In our *treeclimbR* package, 25 values of $t$ in the range $[0, 1]$ are chosen from our experience.

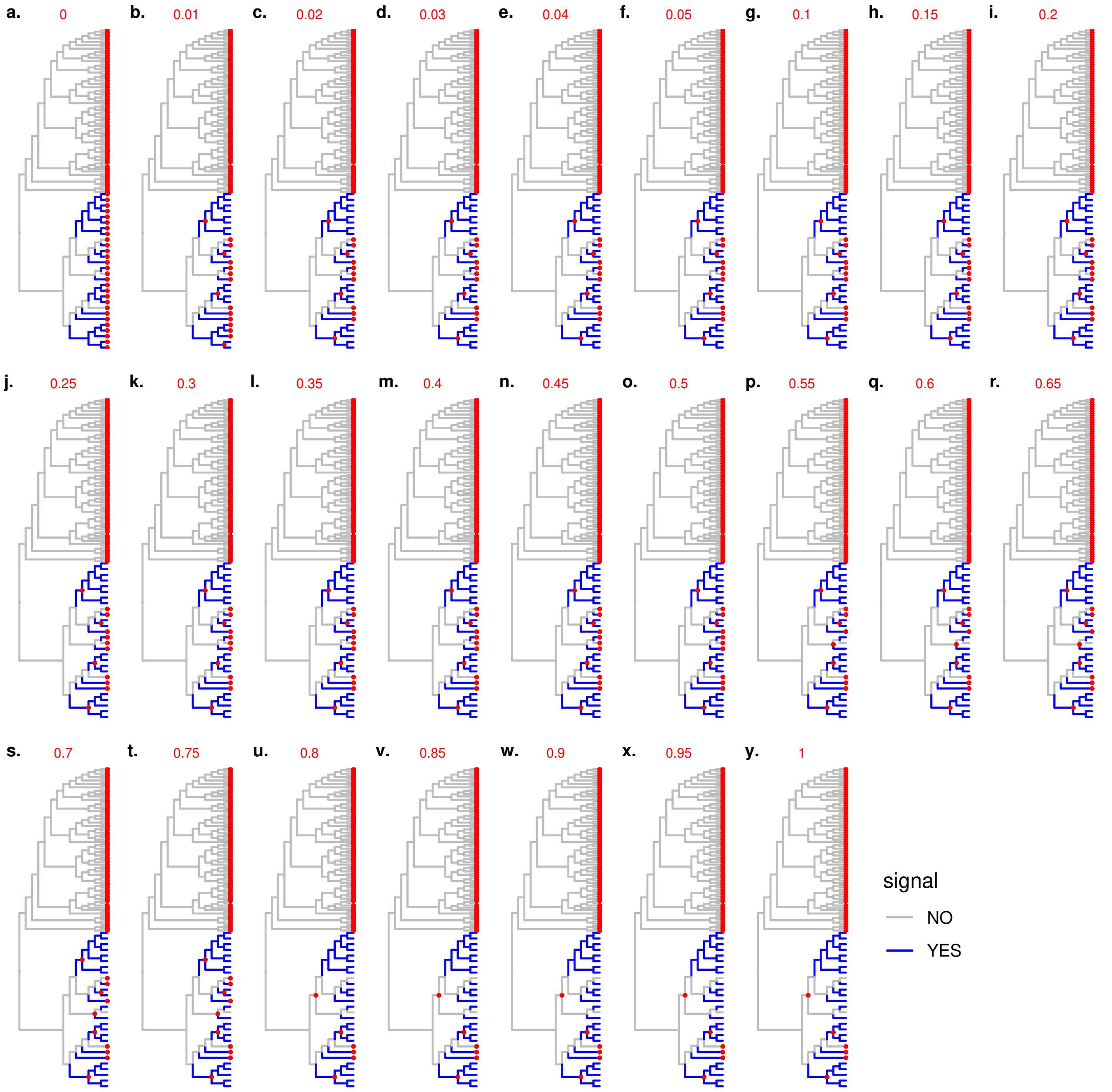

**Supplementary Figure 8. Candidates generated by *treeclimbR* at different $t$ values.** DA branches are colored in blue; otherwise in grey. Nodes that are picked to form a candidate are in red points. The $t$ value used for each candidate is given in the title.

The toy data includes a randomly generated tree with 100 leaves and a count matrix with 100 rows and 40 columns. Each row of the count matrix is matched to a leaf. Each column of the count matrix represents a sample. Counts of leaves within a sample are randomly generated from a multinomial distribution with the total count 1000. The probabilities of leaves are generated randomly from a uniform distribution and scaled to have the sum equal to 1. The first 20 samples (or columns) are labeled as the first group, and the other 20 samples as the second group. We further multiply counts of 25 randomly selected leaves by 3 in the second group. In other words, only 25 leaves are different between groups.

*treeclimbR* is applied on this toy data, and candidates generated at the default values of $t$ in our *treeclimbR* packages are shown in Supplementary Figure 8. When $t = 0$, the candidate is on the leaf level. As $t$ increases, multiple DA leaves are represented by internal nodes. When $t = 0.02$, the signal pattern on the tree is captured perfectly. Even when $t$ is further increased, the candidate stays in the same level until $t = 0.3$. In non-DA branches, leaves are picked even at $t = 1$.

### Supplementary Note 3: The *minP* procedure

*minP* was our initial idea to identify signal branches. It includes three main steps: data aggregation, differential analysis, and node selection. The first two steps are the same as in *treeclimbR*. Data on internal nodes are generated from that on leaves, and differential analysis is performed on all nodes of the tree for each feature as

$$\mathbf{p}^g = (p_1^g, p_1^g, ..., p_M^g); \quad g \in (1, 2, ..., G) \tag{27}$$

where $M$ is the total number of nodes on the tree and $g$ represents a feature. Hypotheses on all nodes and features are pooled, and multiplicity correction is performed using the *BH* [3] procedure. Nodes with null hypotheses rejected are collected as a set, $S_1$. The final step is based on P-values.

1. It starts from the root and moves toward leaves along edges.
2. For each path, it stops when a node $i$ has $p_i \leq p_k$ where node $k$ belongs to descendants of node $i$ ($k \in D(i)$). Nodes that are termini of paths are collected as a set, $S_2$.
3. Finally, nodes in both $S_1$ and $S_2$ are selected.

In other words, a node is selected by *minP* only when its null hypothesis is rejected and none of its descendants have lower P-value.
