## Supplementary (countSimQC report) for "treeclimbR pinpoints the data-dependent resolution of hierarchical hypotheses"

Comparison of characteristic features across count data sets


Code 

- Show All Code
- Hide All Code

### Comparison of characteristic features across count data sets

#### Description

This is a comparison between real and simulated microbial data on throat

###### Design formulae

The following model formulae were used in the dispersion calculations for the different data sets. Note that if a count matrix or data frame was provided in place of a *DESeqDataSet* for some data set, the corresponding design formula is set to `~1`, thus assuming that all samples are to be treated as replicates.

```
original :  ~ 1 
simulate :  ~ 1
```

#### Data set dimensions

These bar plots show the number of samples (columns) and features (rows) in each data set.

##### Number of samples (columns)

##### Number of features (rows)

#### Dispersion/BCV plots

Disperson/BCV plots show the association between the average abundance and the dispersion or “biological coefficient of variation” (sqrt(dispersion)), as calculated by `edgeR` (Robinson, McCarthy, and Smyth 2010) and `DESeq2` (Love, Huber, and Anders 2014). In the `edgeR` plot, the estimate of the prior degrees of freedom is indicated.

##### edgeR

The black dots represent the tagwise dispersion estimates, the red line the common dispersion and the blue curve represents the trended dispersion estimates. For further information about the dispersion estimation in `edgeR`, see Chen, Lun, and Smyth (2014).

##### Pairwise comparisons - edgeR

The table below contains a range of quantitative statistics and test results evaluating the degree of similarity between each pair of data sets based on the average log CPM and tagwise dispersion. The following statistics and test results are included:

- *NN rejection fraction*: this statistic is similar to one used by the kBET R package. First, a subset of R features are selected (where R is chosen to be the smallest of 500 and the total number of features in the two data sets), and for each of these features the k nearest neighbors are found (where k is chosen to be 1% of the total number of features from the two data sets, however at least 5). Then, a chi-square test is used to evaluate whether the composition of data sets from which the k nearest neighbors of each selected features comes differs significantly from the overall composition of features from the two data sets. The NN rejection fraction is the fraction of selected features for which the chi-square test rejects the null hypothesis at a significance level of 5%.
- *Average silhouette width*: as for the NN rejection fraction, a subset of R features are selected. For each of these, the (Euclidean) distances to all other features are calculated, and the silhouette width (Rousseeuw 1987) is calculated based on these distances. The final statistic is the average silhouette width for the R features.
- *Average local silhouette width*: similar to the average silhouette width, but for each feature, only the k’ closest features from a data set are used to calculate the silhouette widths. Here, k’ for data set i is defined as k \* p\_i, where k is as above and p\_i is the fraction of the total number of features that come from data set i.
- For the NN rejection fraction and average silhouette widths, permutation p-values can be calculated by permuting the data set labels. These values are only available if the ‘permutationPvalues’ argument to ‘countsimQCReport’ was set to TRUE.

##### DESeq2

The black dots are the gene-wise dispersion estimates, the red curve the fitted mean-dispersion relationship and the blue circles represent the final dispersion estimates.For further information about the dispersion estimation in `DESeq2`, see Love, Huber, and Anders (2014).

##### Pairwise comparisons - DESeq2

The table below contains a range of quantitative statistics and test results evaluating the degree of similarity between each pair of data sets based on the base mean and genewise dispersion. The following statistics and test results are included:

- *NN rejection fraction*: this statistic is similar to one used by the kBET R package. First, a subset of R features are selected (where R is chosen to be the smallest of 500 and the total number of features in the two data sets), and for each of these features the k nearest neighbors are found (where k is chosen to be 1% of the total number of features from the two data sets, however at least 5). Then, a chi-square test is used to evaluate whether the composition of data sets from which the k nearest neighbors of each selected features comes differs significantly from the overall composition of features from the two data sets. The NN rejection fraction is the fraction of selected features for which the chi-square test rejects the null hypothesis at a significance level of 5%.
- *Average silhouette width*: as for the NN rejection fraction, a subset of R features are selected. For each of these, the (Euclidean) distances to all other features are calculated, and the silhouette width (Rousseeuw 1987) is calculated based on these distances. The final statistic is the average silhouette width for the R features.
- *Average local silhouette width*: similar to the average silhouette width, but for each feature, only the k’ closest features from a data set are used to calculate the silhouette widths. Here, k’ for data set i is defined as k \* p\_i, where k is as above and p\_i is the fraction of the total number of features that come from data set i.
- For the NN rejection fraction and average silhouette widths, permutation p-values can be calculated by permuting the data set labels. These values are only available if the ‘permutationPvalues’ argument to ‘countsimQCReport’ was set to TRUE.

#### Mean-variance plots

These scatter plots show the relation between the empirical mean and variance of the features. The difference between these mean-variance plots and the mean-dispersion plots above is that the plots in this section do not take the information about the experimental design and sample grouping into account, but simply display the mean and variance of log2(CPM) estimates across all samples, calculated using the `cpm` function from `edgeR` (Robinson, McCarthy, and Smyth 2010), with a prior count of 2.

##### Separate scatter plots

##### Overlaid scatter plots

##### Pairwise comparisons

The table below contains a range of quantitative statistics and test results evaluating the degree of similarity between each pair of data sets based on the average and variance of the log CPM. The following statistics and test results are included:

- *NN rejection fraction*: this statistic is similar to one used by the kBET R package. First, a subset of R features are selected (where R is chosen to be the smallest of 500 and the total number of features in the two data sets), and for each of these features the k nearest neighbors are found (where k is chosen to be 1% of the total number of features from the two data sets, however at least 5). Then, a chi-square test is used to evaluate whether the composition of data sets from which the k nearest neighbors of each selected features comes differs significantly from the overall composition of features from the two data sets. The NN rejection fraction is the fraction of selected features for which the chi-square test rejects the null hypothesis at a significance level of 5%.
- *Average silhouette width*: as for the NN rejection fraction, a subset of R features are selected. For each of these, the (Euclidean) distances to all other features are calculated, and the silhouette width (Rousseeuw 1987) is calculated based on these distances. The final statistic is the average silhouette width for the R features.
- *Average local silhouette width*: similar to the average silhouette width, but for each feature, only the k’ closest features from a data set are used to calculate the silhouette widths. Here, k’ for data set i is defined as k \* p\_i, where k is as above and p\_i is the fraction of the total number of features that come from data set i.
- For the NN rejection fraction and average silhouette widths, permutation p-values can be calculated by permuting the data set labels. These values are only available if the ‘permutationPvalues’ argument to ‘countsimQCReport’ was set to TRUE.

#### Library sizes

These plots illustrate the distribution of the total read count per sample, i.e., the column sums of the respective data matrices.

##### Separate histograms

##### Overlaid histograms

##### Density plots

##### Density plots (filled)

##### Box plots

##### Violin plots

##### Empirical cumulative distribution function

##### Pairwise comparisons

The table below contains a range of quantitative statistics and test results evaluating the degree of similarity between each pair of data sets based on the library size. The following statistics and test results are included:

- *K-S statistic/K-S p-value*: the Kolmogorov-Smirnov statistic (Kolmogorov 1933, @Smirnov1948), i.e., the maximal distance between the two empirical cumulative distribution functions (eCDFs), and the associated p-value.
- *Scaled area between eCDFs*: the absolute area between the eCDFs for the two data sets. This is calculated by first subtracting one eCDF from the other and taking the absolute value of the differences, followed by scaling the x-axis so that the difference between the largest and smallest value across all data sets is equal to 1 (here, subtracting 1278 and dividing by 4133), and finally calculating the area under the resulting non-negative curve. This gives a scaled area with values in the range [0, 1], where large values indicate large differences between the library size distributions.
- *Runs statistic/Runs p-value*: the statistic and p-value from a one-sided Wald-Wolfowitz runs test (Wald and Wolfowitz 1940). The values from both data sets are first sorted in increasing order and converted into a binary sequence based on whether or not they are derived from the first data set. The runs test is applied to this sequence. The test is one-sided, with the alternative hypothesis being that there is a trend towards values from the same data set occurring next to each other in the sequence.
- *NN rejection fraction*: this statistic is similar to one used by the kBET R package. First, a subset of R samples are selected (where R is chosen to be the smallest of 500 and the total number of samples in the two data sets), and for each of these samples the k nearest neighbors are found (where k is chosen to be 1% of the total number of samples from the two data sets, however at least 5). Then, a chi-square test is used to evaluate whether the composition of data sets from which the k nearest neighbors of each selected samples comes differs significantly from the overall composition of samples from the two data sets. The NN rejection fraction is the fraction of selected samples for which the chi-square test rejects the null hypothesis at a significance level of 5%.
- *Average silhouette width*: as for the NN rejection fraction, a subset of R samples are selected. For each of these, the (Euclidean) distances to all other samples are calculated, and the silhouette width (Rousseeuw 1987) is calculated based on these distances. The final statistic is the average silhouette width for the R samples.
- *Average local silhouette width*: similar to the average silhouette width, but for each sample, only the k’ closest samples from a data set are used to calculate the silhouette widths. Here, k’ for data set i is defined as k \* p\_i, where k is as above and p\_i is the fraction of the total number of samples that come from data set i.
- For the scaled area between eCDFs, NN rejection fraction and average silhouette widths, permutation p-values can be calculated by permuting the data set labels. These values are only available if the ‘permutationPvalues’ argument to ‘countsimQCReport’ was set to TRUE.

It should be noted that the reported statistics in some cases are highly dependent on the number of underlying observations, and especially with a large number of observations, very small p-values can correspond to small effective differences. Thus, interpretation should ideally be guided by both quantitative and qualitative, visual information.

#### TMM normalization factors

The plots below show the distribution of the TMM normalization factors (Robinson and Oshlack 2010), intended to adjust for differences in RNA composition, as calculated by `edgeR` (Robinson, McCarthy, and Smyth 2010).

##### Separate histograms

##### Overlaid histograms

##### Density plots

##### Density plots (filled)

##### Box plots

##### Violin plots

##### Empirical cumulative distribution function

##### Pairwise comparisons

The table below contains a range of quantitative statistics and test results evaluating the degree of similarity between each pair of data sets based on the TMM factor. The following statistics and test results are included:

- *K-S statistic/K-S p-value*: the Kolmogorov-Smirnov statistic (Kolmogorov 1933, @Smirnov1948), i.e., the maximal distance between the two empirical cumulative distribution functions (eCDFs), and the associated p-value.
- *Scaled area between eCDFs*: the absolute area between the eCDFs for the two data sets. This is calculated by first subtracting one eCDF from the other and taking the absolute value of the differences, followed by scaling the x-axis so that the difference between the largest and smallest value across all data sets is equal to 1 (here, subtracting 0.365578817943224 and dividing by 2.94720337367015), and finally calculating the area under the resulting non-negative curve. This gives a scaled area with values in the range [0, 1], where large values indicate large differences between the TMM factor distributions.
- *Runs statistic/Runs p-value*: the statistic and p-value from a one-sided Wald-Wolfowitz runs test (Wald and Wolfowitz 1940). The values from both data sets are first sorted in increasing order and converted into a binary sequence based on whether or not they are derived from the first data set. The runs test is applied to this sequence. The test is one-sided, with the alternative hypothesis being that there is a trend towards values from the same data set occurring next to each other in the sequence.
- *NN rejection fraction*: this statistic is similar to one used by the kBET R package. First, a subset of R samples are selected (where R is chosen to be the smallest of 500 and the total number of samples in the two data sets), and for each of these samples the k nearest neighbors are found (where k is chosen to be 1% of the total number of samples from the two data sets, however at least 5). Then, a chi-square test is used to evaluate whether the composition of data sets from which the k nearest neighbors of each selected samples comes differs significantly from the overall composition of samples from the two data sets. The NN rejection fraction is the fraction of selected samples for which the chi-square test rejects the null hypothesis at a significance level of 5%.
- *Average silhouette width*: as for the NN rejection fraction, a subset of R samples are selected. For each of these, the (Euclidean) distances to all other samples are calculated, and the silhouette width (Rousseeuw 1987) is calculated based on these distances. The final statistic is the average silhouette width for the R samples.
- *Average local silhouette width*: similar to the average silhouette width, but for each sample, only the k’ closest samples from a data set are used to calculate the silhouette widths. Here, k’ for data set i is defined as k \* p\_i, where k is as above and p\_i is the fraction of the total number of samples that come from data set i.
- For the scaled area between eCDFs, NN rejection fraction and average silhouette widths, permutation p-values can be calculated by permuting the data set labels. These values are only available if the ‘permutationPvalues’ argument to ‘countsimQCReport’ was set to TRUE.

It should be noted that the reported statistics in some cases are highly dependent on the number of underlying observations, and especially with a large number of observations, very small p-values can correspond to small effective differences. Thus, interpretation should ideally be guided by both quantitative and qualitative, visual information.

#### Effective library sizes

These plots show the distribution of the “effective library sizes”, defined as the total count per sample multiplied by the corresponding TMM normalization factor.

##### Separate histograms

##### Overlaid histograms

##### Density plots

##### Density plots (filled)

##### Box plots

##### Violin plots

##### Empirical cumulative distribution function

##### Pairwise comparisons

The table below contains a range of quantitative statistics and test results evaluating the degree of similarity between each pair of data sets based on the effective library size. The following statistics and test results are included:

- *K-S statistic/K-S p-value*: the Kolmogorov-Smirnov statistic (Kolmogorov 1933, @Smirnov1948), i.e., the maximal distance between the two empirical cumulative distribution functions (eCDFs), and the associated p-value.
- *Scaled area between eCDFs*: the absolute area between the eCDFs for the two data sets. This is calculated by first subtracting one eCDF from the other and taking the absolute value of the differences, followed by scaling the x-axis so that the difference between the largest and smallest value across all data sets is equal to 1 (here, subtracting 1465.50757879645 and dividing by 4341.7996031018), and finally calculating the area under the resulting non-negative curve. This gives a scaled area with values in the range [0, 1], where large values indicate large differences between the effective library size distributions.
- *Runs statistic/Runs p-value*: the statistic and p-value from a one-sided Wald-Wolfowitz runs test (Wald and Wolfowitz 1940). The values from both data sets are first sorted in increasing order and converted into a binary sequence based on whether or not they are derived from the first data set. The runs test is applied to this sequence. The test is one-sided, with the alternative hypothesis being that there is a trend towards values from the same data set occurring next to each other in the sequence.
- *NN rejection fraction*: this statistic is similar to one used by the kBET R package. First, a subset of R samples are selected (where R is chosen to be the smallest of 500 and the total number of samples in the two data sets), and for each of these samples the k nearest neighbors are found (where k is chosen to be 1% of the total number of samples from the two data sets, however at least 5). Then, a chi-square test is used to evaluate whether the composition of data sets from which the k nearest neighbors of each selected samples comes differs significantly from the overall composition of samples from the two data sets. The NN rejection fraction is the fraction of selected samples for which the chi-square test rejects the null hypothesis at a significance level of 5%.
- *Average silhouette width*: as for the NN rejection fraction, a subset of R samples are selected. For each of these, the (Euclidean) distances to all other samples are calculated, and the silhouette width (Rousseeuw 1987) is calculated based on these distances. The final statistic is the average silhouette width for the R samples.
- *Average local silhouette width*: similar to the average silhouette width, but for each sample, only the k’ closest samples from a data set are used to calculate the silhouette widths. Here, k’ for data set i is defined as k \* p\_i, where k is as above and p\_i is the fraction of the total number of samples that come from data set i.
- For the scaled area between eCDFs, NN rejection fraction and average silhouette widths, permutation p-values can be calculated by permuting the data set labels. These values are only available if the ‘permutationPvalues’ argument to ‘countsimQCReport’ was set to TRUE.

It should be noted that the reported statistics in some cases are highly dependent on the number of underlying observations, and especially with a large number of observations, very small p-values can correspond to small effective differences. Thus, interpretation should ideally be guided by both quantitative and qualitative, visual information.

#### Expression distributions (average log CPM)

The plots in this section show the distribution of average abundance values for the features. The abundances are log CPM values calculated by `edgeR`.

##### Separate histograms

##### Overlaid histograms

##### Density plots

##### Density plots (filled)

##### Box plots

##### Violin plots

##### Empirical cumulative distribution function

##### Pairwise comparisons

The table below contains a range of quantitative statistics and test results evaluating the degree of similarity between each pair of data sets based on the average log CPM. The following statistics and test results are included:

- *K-S statistic/K-S p-value*: the Kolmogorov-Smirnov statistic (Kolmogorov 1933, @Smirnov1948), i.e., the maximal distance between the two empirical cumulative distribution functions (eCDFs), and the associated p-value.
- *Scaled area between eCDFs*: the absolute area between the eCDFs for the two data sets. This is calculated by first subtracting one eCDF from the other and taking the absolute value of the differences, followed by scaling the x-axis so that the difference between the largest and smallest value across all data sets is equal to 1 (here, subtracting 9.41981795160182 and dividing by 6.58462849082064), and finally calculating the area under the resulting non-negative curve. This gives a scaled area with values in the range [0, 1], where large values indicate large differences between the average log CPM distributions.
- *Runs statistic/Runs p-value*: the statistic and p-value from a one-sided Wald-Wolfowitz runs test (Wald and Wolfowitz 1940). The values from both data sets are first sorted in increasing order and converted into a binary sequence based on whether or not they are derived from the first data set. The runs test is applied to this sequence. The test is one-sided, with the alternative hypothesis being that there is a trend towards values from the same data set occurring next to each other in the sequence.
- *NN rejection fraction*: this statistic is similar to one used by the kBET R package. First, a subset of R features are selected (where R is chosen to be the smallest of 500 and the total number of features in the two data sets), and for each of these features the k nearest neighbors are found (where k is chosen to be 1% of the total number of features from the two data sets, however at least 5). Then, a chi-square test is used to evaluate whether the composition of data sets from which the k nearest neighbors of each selected features comes differs significantly from the overall composition of features from the two data sets. The NN rejection fraction is the fraction of selected features for which the chi-square test rejects the null hypothesis at a significance level of 5%.
- *Average silhouette width*: as for the NN rejection fraction, a subset of R features are selected. For each of these, the (Euclidean) distances to all other features are calculated, and the silhouette width (Rousseeuw 1987) is calculated based on these distances. The final statistic is the average silhouette width for the R features.
- *Average local silhouette width*: similar to the average silhouette width, but for each feature, only the k’ closest features from a data set are used to calculate the silhouette widths. Here, k’ for data set i is defined as k \* p\_i, where k is as above and p\_i is the fraction of the total number of features that come from data set i.
- For the scaled area between eCDFs, NN rejection fraction and average silhouette widths, permutation p-values can be calculated by permuting the data set labels. These values are only available if the ‘permutationPvalues’ argument to ‘countsimQCReport’ was set to TRUE.

It should be noted that the reported statistics in some cases are highly dependent on the number of underlying observations, and especially with a large number of observations, very small p-values can correspond to small effective differences. Thus, interpretation should ideally be guided by both quantitative and qualitative, visual information.

#### Fraction zeros per sample

These plots show the distribution of the fraction of zeros observed per sample (column) in the count matrices.

##### Separate histograms

##### Overlaid histograms

##### Density plots

##### Density plots (filled)

##### Box plots

##### Violin plots

##### Empirical cumulative distribution function

##### Pairwise comparisons

The table below contains a range of quantitative statistics and test results evaluating the degree of similarity between each pair of data sets based on the fraction zeros. The following statistics and test results are included:

- *K-S statistic/K-S p-value*: the Kolmogorov-Smirnov statistic (Kolmogorov 1933, @Smirnov1948), i.e., the maximal distance between the two empirical cumulative distribution functions (eCDFs), and the associated p-value.
- *Scaled area between eCDFs*: the absolute area between the eCDFs for the two data sets. This is calculated by first subtracting one eCDF from the other and taking the absolute value of the differences, followed by scaling the x-axis so that the difference between the largest and smallest value across all data sets is equal to 1 (here, subtracting 0.383891213389121 and dividing by 0.494769874476987), and finally calculating the area under the resulting non-negative curve. This gives a scaled area with values in the range [0, 1], where large values indicate large differences between the fraction zeros distributions.
- *Runs statistic/Runs p-value*: the statistic and p-value from a one-sided Wald-Wolfowitz runs test (Wald and Wolfowitz 1940). The values from both data sets are first sorted in increasing order and converted into a binary sequence based on whether or not they are derived from the first data set. The runs test is applied to this sequence. The test is one-sided, with the alternative hypothesis being that there is a trend towards values from the same data set occurring next to each other in the sequence.
- *NN rejection fraction*: this statistic is similar to one used by the kBET R package. First, a subset of R samples are selected (where R is chosen to be the smallest of 500 and the total number of samples in the two data sets), and for each of these samples the k nearest neighbors are found (where k is chosen to be 1% of the total number of samples from the two data sets, however at least 5). Then, a chi-square test is used to evaluate whether the composition of data sets from which the k nearest neighbors of each selected samples comes differs significantly from the overall composition of samples from the two data sets. The NN rejection fraction is the fraction of selected samples for which the chi-square test rejects the null hypothesis at a significance level of 5%.
- *Average silhouette width*: as for the NN rejection fraction, a subset of R samples are selected. For each of these, the (Euclidean) distances to all other samples are calculated, and the silhouette width (Rousseeuw 1987) is calculated based on these distances. The final statistic is the average silhouette width for the R samples.
- *Average local silhouette width*: similar to the average silhouette width, but for each sample, only the k’ closest samples from a data set are used to calculate the silhouette widths. Here, k’ for data set i is defined as k \* p\_i, where k is as above and p\_i is the fraction of the total number of samples that come from data set i.
- For the scaled area between eCDFs, NN rejection fraction and average silhouette widths, permutation p-values can be calculated by permuting the data set labels. These values are only available if the ‘permutationPvalues’ argument to ‘countsimQCReport’ was set to TRUE.

It should be noted that the reported statistics in some cases are highly dependent on the number of underlying observations, and especially with a large number of observations, very small p-values can correspond to small effective differences. Thus, interpretation should ideally be guided by both quantitative and qualitative, visual information.

#### Fraction zeros per feature

These plots illustrate the distribution of the fraction of zeros observed per feature (row) in the count matrices.

##### Separate histograms

##### Overlaid histograms

##### Density plots

##### Density plots (filled)

##### Box plots

##### Violin plots

##### Empirical cumulative distribution function

##### Pairwise comparisons

The table below contains a range of quantitative statistics and test results evaluating the degree of similarity between each pair of data sets based on the fraction zeros. The following statistics and test results are included:

- *K-S statistic/K-S p-value*: the Kolmogorov-Smirnov statistic (Kolmogorov 1933, @Smirnov1948), i.e., the maximal distance between the two empirical cumulative distribution functions (eCDFs), and the associated p-value.
- *Scaled area between eCDFs*: the absolute area between the eCDFs for the two data sets. This is calculated by first subtracting one eCDF from the other and taking the absolute value of the differences, followed by scaling the x-axis so that the difference between the largest and smallest value across all data sets is equal to 1 (here, subtracting 0 and dividing by 0.88), and finally calculating the area under the resulting non-negative curve. This gives a scaled area with values in the range [0, 1], where large values indicate large differences between the fraction zeros distributions.
- *Runs statistic/Runs p-value*: the statistic and p-value from a one-sided Wald-Wolfowitz runs test (Wald and Wolfowitz 1940). The values from both data sets are first sorted in increasing order and converted into a binary sequence based on whether or not they are derived from the first data set. The runs test is applied to this sequence. The test is one-sided, with the alternative hypothesis being that there is a trend towards values from the same data set occurring next to each other in the sequence.
- *NN rejection fraction*: this statistic is similar to one used by the kBET R package. First, a subset of R features are selected (where R is chosen to be the smallest of 500 and the total number of features in the two data sets), and for each of these features the k nearest neighbors are found (where k is chosen to be 1% of the total number of features from the two data sets, however at least 5). Then, a chi-square test is used to evaluate whether the composition of data sets from which the k nearest neighbors of each selected features comes differs significantly from the overall composition of features from the two data sets. The NN rejection fraction is the fraction of selected features for which the chi-square test rejects the null hypothesis at a significance level of 5%.
- *Average silhouette width*: as for the NN rejection fraction, a subset of R features are selected. For each of these, the (Euclidean) distances to all other features are calculated, and the silhouette width (Rousseeuw 1987) is calculated based on these distances. The final statistic is the average silhouette width for the R features.
- *Average local silhouette width*: similar to the average silhouette width, but for each feature, only the k’ closest features from a data set are used to calculate the silhouette widths. Here, k’ for data set i is defined as k \* p\_i, where k is as above and p\_i is the fraction of the total number of features that come from data set i.
- For the scaled area between eCDFs, NN rejection fraction and average silhouette widths, permutation p-values can be calculated by permuting the data set labels. These values are only available if the ‘permutationPvalues’ argument to ‘countsimQCReport’ was set to TRUE.

It should be noted that the reported statistics in some cases are highly dependent on the number of underlying observations, and especially with a large number of observations, very small p-values can correspond to small effective differences. Thus, interpretation should ideally be guided by both quantitative and qualitative, visual information.

#### Sample-sample correlations

The plots below show the distribution of Spearman correlation coefficients for pairs of samples, calculated from the log(CPM) values obtained via the `cpm` function from `edgeR`, with a prior.count of 2. If there are more than 500 samples in a data set, the pairwise correlations between 500 randomly selected samples are shown.

##### Separate histograms

##### Overlaid histograms

##### Density plots

##### Density plots (filled)

##### Box plots

##### Violin plots

##### Empirical cumulative distribution function

##### Pairwise comparisons

The table below contains a range of quantitative statistics and test results evaluating the degree of similarity between each pair of data sets based on the Spearman correlation. The following statistics and test results are included:

- *K-S statistic/K-S p-value*: the Kolmogorov-Smirnov statistic (Kolmogorov 1933, @Smirnov1948), i.e., the maximal distance between the two empirical cumulative distribution functions (eCDFs), and the associated p-value.
- *Scaled area between eCDFs*: the absolute area between the eCDFs for the two data sets. This is calculated by first subtracting one eCDF from the other and taking the absolute value of the differences, followed by scaling the x-axis so that the difference between the largest and smallest value across all data sets is equal to 1 (here, subtracting -0.242071291926404 and dividing by 0.928632310646341), and finally calculating the area under the resulting non-negative curve. This gives a scaled area with values in the range [0, 1], where large values indicate large differences between the Spearman correlation distributions.
- *Runs statistic/Runs p-value*: the statistic and p-value from a one-sided Wald-Wolfowitz runs test (Wald and Wolfowitz 1940). The values from both data sets are first sorted in increasing order and converted into a binary sequence based on whether or not they are derived from the first data set. The runs test is applied to this sequence. The test is one-sided, with the alternative hypothesis being that there is a trend towards values from the same data set occurring next to each other in the sequence.
- *NN rejection fraction*: this statistic is similar to one used by the kBET R package. First, a subset of R sample pairs are selected (where R is chosen to be the smallest of 500 and the total number of sample pairs in the two data sets), and for each of these sample pairs the k nearest neighbors are found (where k is chosen to be 1% of the total number of sample pairs from the two data sets, however at least 5). Then, a chi-square test is used to evaluate whether the composition of data sets from which the k nearest neighbors of each selected sample pairs comes differs significantly from the overall composition of sample pairs from the two data sets. The NN rejection fraction is the fraction of selected sample pairs for which the chi-square test rejects the null hypothesis at a significance level of 5%.
- *Average silhouette width*: as for the NN rejection fraction, a subset of R sample pairs are selected. For each of these, the (Euclidean) distances to all other sample pairs are calculated, and the silhouette width (Rousseeuw 1987) is calculated based on these distances. The final statistic is the average silhouette width for the R sample pairs.
- *Average local silhouette width*: similar to the average silhouette width, but for each sample pair, only the k’ closest sample pairs from a data set are used to calculate the silhouette widths. Here, k’ for data set i is defined as k \* p\_i, where k is as above and p\_i is the fraction of the total number of sample pairs that come from data set i.
- For the scaled area between eCDFs, NN rejection fraction and average silhouette widths, permutation p-values can be calculated by permuting the data set labels. These values are only available if the ‘permutationPvalues’ argument to ‘countsimQCReport’ was set to TRUE.

It should be noted that the reported statistics in some cases are highly dependent on the number of underlying observations, and especially with a large number of observations, very small p-values can correspond to small effective differences. Thus, interpretation should ideally be guided by both quantitative and qualitative, visual information.

#### Feature-feature correlations

These plots illustrate the distribution of Spearman correlation coefficients for pairs of features, calculated from the log(CPM) values obtained via the `cpm` function from `edgeR`, with a prior.count of 2. Only non-constant features are considered, and if there are more than 500 such features in a data set, the pairwise correlations between 500 randomly selected features are shown.

##### Separate histograms

##### Overlaid histograms

##### Density plots

##### Density plots (filled)

##### Box plots

##### Violin plots

##### Empirical cumulative distribution function

##### Pairwise comparisons

The table below contains a range of quantitative statistics and test results evaluating the degree of similarity between each pair of data sets based on the Spearman correlation. The following statistics and test results are included:

- *K-S statistic/K-S p-value*: the Kolmogorov-Smirnov statistic (Kolmogorov 1933, @Smirnov1948), i.e., the maximal distance between the two empirical cumulative distribution functions (eCDFs), and the associated p-value.
- *Scaled area between eCDFs*: the absolute area between the eCDFs for the two data sets. This is calculated by first subtracting one eCDF from the other and taking the absolute value of the differences, followed by scaling the x-axis so that the difference between the largest and smallest value across all data sets is equal to 1 (here, subtracting -0.671556511984037 and dividing by 1.51034432692276), and finally calculating the area under the resulting non-negative curve. This gives a scaled area with values in the range [0, 1], where large values indicate large differences between the Spearman correlation distributions.
- *Runs statistic/Runs p-value*: the statistic and p-value from a one-sided Wald-Wolfowitz runs test (Wald and Wolfowitz 1940). The values from both data sets are first sorted in increasing order and converted into a binary sequence based on whether or not they are derived from the first data set. The runs test is applied to this sequence. The test is one-sided, with the alternative hypothesis being that there is a trend towards values from the same data set occurring next to each other in the sequence.
- *NN rejection fraction*: this statistic is similar to one used by the kBET R package. First, a subset of R feature pairs are selected (where R is chosen to be the smallest of 500 and the total number of feature pairs in the two data sets), and for each of these feature pairs the k nearest neighbors are found (where k is chosen to be 1% of the total number of feature pairs from the two data sets, however at least 5). Then, a chi-square test is used to evaluate whether the composition of data sets from which the k nearest neighbors of each selected feature pairs comes differs significantly from the overall composition of feature pairs from the two data sets. The NN rejection fraction is the fraction of selected feature pairs for which the chi-square test rejects the null hypothesis at a significance level of 5%.
- *Average silhouette width*: as for the NN rejection fraction, a subset of R feature pairs are selected. For each of these, the (Euclidean) distances to all other feature pairs are calculated, and the silhouette width (Rousseeuw 1987) is calculated based on these distances. The final statistic is the average silhouette width for the R feature pairs.
- *Average local silhouette width*: similar to the average silhouette width, but for each feature pair, only the k’ closest feature pairs from a data set are used to calculate the silhouette widths. Here, k’ for data set i is defined as k \* p\_i, where k is as above and p\_i is the fraction of the total number of feature pairs that come from data set i.
- For the scaled area between eCDFs, NN rejection fraction and average silhouette widths, permutation p-values can be calculated by permuting the data set labels. These values are only available if the ‘permutationPvalues’ argument to ‘countsimQCReport’ was set to TRUE.

It should be noted that the reported statistics in some cases are highly dependent on the number of underlying observations, and especially with a large number of observations, very small p-values can correspond to small effective differences. Thus, interpretation should ideally be guided by both quantitative and qualitative, visual information.

#### Library size vs fraction zeros

These scatter plots show the association between the total count (column sums) and the fraction of zeros observed per sample.

##### Separate scatter plots

##### Overlaid scatter plots

##### Pairwise comparisons

The table below contains a range of quantitative statistics and test results evaluating the degree of similarity between each pair of data sets based on the library size and fraction of zeros. The following statistics and test results are included:

- *NN rejection fraction*: this statistic is similar to one used by the kBET R package. First, a subset of R samples are selected (where R is chosen to be the smallest of 500 and the total number of samples in the two data sets), and for each of these samples the k nearest neighbors are found (where k is chosen to be 1% of the total number of samples from the two data sets, however at least 5). Then, a chi-square test is used to evaluate whether the composition of data sets from which the k nearest neighbors of each selected samples comes differs significantly from the overall composition of samples from the two data sets. The NN rejection fraction is the fraction of selected samples for which the chi-square test rejects the null hypothesis at a significance level of 5%.
- *Average silhouette width*: as for the NN rejection fraction, a subset of R samples are selected. For each of these, the (Euclidean) distances to all other samples are calculated, and the silhouette width (Rousseeuw 1987) is calculated based on these distances. The final statistic is the average silhouette width for the R samples.
- *Average local silhouette width*: similar to the average silhouette width, but for each sample, only the k’ closest samples from a data set are used to calculate the silhouette widths. Here, k’ for data set i is defined as k \* p\_i, where k is as above and p\_i is the fraction of the total number of samples that come from data set i.
- For the NN rejection fraction and average silhouette widths, permutation p-values can be calculated by permuting the data set labels. These values are only available if the ‘permutationPvalues’ argument to ‘countsimQCReport’ was set to TRUE.

#### Mean expression vs fraction zeros

These scatter plots show the association between the average abundance and the fraction of zeros observed per feature. The abundance is defined as the log(CPM) values as calculated by `edgeR`.

##### Separate scatter plots

##### Overlaid scatter plots

##### Pairwise comparisons

The table below contains a range of quantitative statistics and test results evaluating the degree of similarity between each pair of data sets based on the average log CPM and fraction of zeros. The following statistics and test results are included:

- *NN rejection fraction*: this statistic is similar to one used by the kBET R package. First, a subset of R features are selected (where R is chosen to be the smallest of 500 and the total number of features in the two data sets), and for each of these features the k nearest neighbors are found (where k is chosen to be 1% of the total number of features from the two data sets, however at least 5). Then, a chi-square test is used to evaluate whether the composition of data sets from which the k nearest neighbors of each selected features comes differs significantly from the overall composition of features from the two data sets. The NN rejection fraction is the fraction of selected features for which the chi-square test rejects the null hypothesis at a significance level of 5%.
- *Average silhouette width*: as for the NN rejection fraction, a subset of R features are selected. For each of these, the (Euclidean) distances to all other features are calculated, and the silhouette width (Rousseeuw 1987) is calculated based on these distances. The final statistic is the average silhouette width for the R features.
- *Average local silhouette width*: similar to the average silhouette width, but for each feature, only the k’ closest features from a data set are used to calculate the silhouette widths. Here, k’ for data set i is defined as k \* p\_i, where k is as above and p\_i is the fraction of the total number of features that come from data set i.
- For the NN rejection fraction and average silhouette widths, permutation p-values can be calculated by permuting the data set labels. These values are only available if the ‘permutationPvalues’ argument to ‘countsimQCReport’ was set to TRUE.

#### Session info

```
## R version 3.6.1 (2019-07-05)
## Platform: x86_64-apple-darwin15.6.0 (64-bit)
## Running under: macOS Mojave 10.14.4
## 
## Matrix products: default
## BLAS:   /System/Library/Frameworks/Accelerate.framework/Versions/A/Frameworks/vecLib.framework/Versions/A/libBLAS.dylib
## LAPACK: /Library/Frameworks/R.framework/Versions/3.6/Resources/lib/libRlapack.dylib
## 
## locale:
## [1] en_US.UTF-8/en_US.UTF-8/en_US.UTF-8/C/en_US.UTF-8/en_US.UTF-8
## 
## attached base packages:
## [1] parallel  stats4    stats     graphics  grDevices
## [6] utils     datasets  methods   base     
## 
## other attached packages:
##  [1] treeclimbR_0.1.1              
##  [2] TreeSummarizedExperiment_1.3.0
##  [3] SingleCellExperiment_1.8.0    
##  [4] countsimQC_1.4.0              
##  [5] DESeq2_1.26.0                 
##  [6] SummarizedExperiment_1.16.0   
##  [7] DelayedArray_0.12.0           
##  [8] BiocParallel_1.20.0           
##  [9] matrixStats_0.55.0            
## [10] Biobase_2.46.0                
## [11] GenomicRanges_1.38.0          
## [12] GenomeInfoDb_1.22.0           
## [13] IRanges_2.20.0                
## [14] S4Vectors_0.24.0              
## [15] BiocGenerics_0.32.0           
## 
## loaded via a namespace (and not attached):
##   [1] backports_1.1.6            
##   [2] circlize_0.4.8             
##   [3] Hmisc_4.3-0                
##   [4] diffcyt_1.6.1              
##   [5] igraph_1.2.4.1             
##   [6] plyr_1.8.5                 
##   [7] ConsensusClusterPlus_1.50.0
##   [8] splines_3.6.1              
##   [9] flowCore_1.52.0            
##  [10] crosstalk_1.0.0            
##  [11] fda_2.4.8                  
##  [12] TH.data_1.0-10             
##  [13] ggplot2_3.3.0              
##  [14] digest_0.6.25              
##  [15] htmltools_0.4.0            
##  [16] fansi_0.4.1                
##  [17] magrittr_1.5               
##  [18] checkmate_1.9.4            
##  [19] memoise_1.1.0              
##  [20] CytoML_1.12.0              
##  [21] cluster_2.1.0              
##  [22] ks_1.11.6                  
##  [23] limma_3.42.0               
##  [24] ComplexHeatmap_2.2.0       
##  [25] annotate_1.64.0            
##  [26] RcppParallel_4.4.4         
##  [27] R.utils_2.9.0              
##  [28] sandwich_2.5-1             
##  [29] flowWorkspace_3.34.0       
##  [30] colorspace_1.4-1           
##  [31] blob_1.2.0                 
##  [32] rrcov_1.4-7                
##  [33] xfun_0.11                  
##  [34] dplyr_0.8.5                
##  [35] crayon_1.3.4               
##  [36] RCurl_1.95-4.12            
##  [37] jsonlite_1.6.1             
##  [38] hexbin_1.28.0              
##  [39] graph_1.64.0               
##  [40] lme4_1.1-21                
##  [41] genefilter_1.68.0          
##  [42] dirmult_0.1.3-4            
##  [43] zoo_1.8-6                  
##  [44] survival_2.44-1.1          
##  [45] ape_5.3                    
##  [46] glue_1.4.0                 
##  [47] flowClust_3.24.0           
##  [48] gtable_0.3.0               
##  [49] zlibbioc_1.32.0            
##  [50] XVector_0.26.0             
##  [51] GetoptLong_0.1.7           
##  [52] ggcyto_1.14.0              
##  [53] IDPmisc_1.1.19             
##  [54] Rgraphviz_2.30.0           
##  [55] shape_1.4.4                
##  [56] DEoptimR_1.0-8             
##  [57] scales_1.1.0               
##  [58] mvtnorm_1.0-11             
##  [59] DBI_1.0.0                  
##  [60] edgeR_3.28.0               
##  [61] Rcpp_1.0.4                 
##  [62] xtable_1.8-4               
##  [63] htmlTable_1.13.2           
##  [64] clue_0.3-57                
##  [65] openCyto_1.24.0            
##  [66] foreign_0.8-71             
##  [67] bit_1.1-14                 
##  [68] mclust_5.4.5               
##  [69] FlowSOM_1.18.0             
##  [70] Formula_1.2-3              
##  [71] tsne_0.1-3                 
##  [72] randtests_1.0              
##  [73] DT_0.10                    
##  [74] htmlwidgets_1.5.1          
##  [75] RColorBrewer_1.1-2         
##  [76] acepack_1.4.1              
##  [77] ellipsis_0.3.0             
##  [78] farver_2.0.3               
##  [79] pkgconfig_2.0.3            
##  [80] XML_3.98-1.20              
##  [81] R.methodsS3_1.7.1          
##  [82] nnet_7.3-12                
##  [83] flowViz_1.50.0             
##  [84] locfit_1.5-9.1             
##  [85] later_1.0.0                
##  [86] labeling_0.3               
##  [87] reshape2_1.4.3             
##  [88] flowStats_3.44.0           
##  [89] tidyselect_1.0.0           
##  [90] rlang_0.4.5                
##  [91] AnnotationDbi_1.48.0       
##  [92] munsell_0.5.0              
##  [93] tools_3.6.1                
##  [94] cli_2.0.2                  
##  [95] RSQLite_2.1.2              
##  [96] fastmap_1.0.1              
##  [97] evaluate_0.14              
##  [98] stringr_1.4.0              
##  [99] yaml_2.2.0                 
## [100] knitr_1.26                 
## [101] bit64_0.9-7                
## [102] robustbase_0.93-5          
## [103] caTools_1.17.1.2           
## [104] purrr_0.3.3                
## [105] RBGL_1.62.1                
## [106] nlme_3.1-142               
## [107] mime_0.7                   
## [108] R.oo_1.23.0                
## [109] compiler_3.6.1             
## [110] rstudioapi_0.11            
## [111] png_0.1-7                  
## [112] tibble_3.0.0               
## [113] geneplotter_1.64.0         
## [114] pcaPP_1.9-73               
## [115] stringi_1.4.6              
## [116] lattice_0.20-38            
## [117] Matrix_1.2-17              
## [118] nloptr_1.2.1               
## [119] vctrs_0.2.4                
## [120] pillar_1.4.3               
## [121] lifecycle_0.2.0            
## [122] GlobalOptions_0.1.1        
## [123] data.table_1.12.6          
## [124] bitops_1.0-6               
## [125] corpcor_1.6.9              
## [126] httpuv_1.5.2               
## [127] R6_2.4.1                   
## [128] latticeExtra_0.6-28        
## [129] promises_1.1.0             
## [130] KernSmooth_2.23-15         
## [131] gridExtra_2.3              
## [132] codetools_0.2-16           
## [133] boot_1.3-23                
## [134] MASS_7.3-51.4              
## [135] gtools_3.8.1               
## [136] assertthat_0.2.1           
## [137] rjson_0.2.20               
## [138] withr_2.1.2                
## [139] mnormt_1.5-5               
## [140] multcomp_1.4-10            
## [141] GenomeInfoDbData_1.2.2     
## [142] mgcv_1.8-28                
## [143] ncdfFlow_2.32.0            
## [144] grid_3.6.1                 
## [145] rpart_4.1-15               
## [146] minqa_1.2.4                
## [147] tidyr_1.0.2                
## [148] rmarkdown_1.17             
## [149] shiny_1.4.0                
## [150] base64enc_0.1-3            
## [151] ellipse_0.4.1
```
